# Warburg-like metabolism coordinates FGF and Wnt signaling in the vertebrate embryo

**DOI:** 10.1101/101451

**Authors:** Masayuki Oginuma, Philippe Moncuquet, Fengzhu Xiong, Edward Karoly, Jérome Chal, Karine Guevorkian, Olivier Pourquié

**Affiliations:** Institut de Génétique et de Biologie Moléculaire et Cellulaire (IGBMC), CNRS (UMR 7104), Inserm U964, Université de Strasbourg, Illkirch. F-67400, France.; Department of Genetics, Harvard Medical School and Department of Pathology, Brigham and Woman’s Hospital. 77 Avenue Louis Pasteur, Boston, Massachusetts 02115, USA.; Metabolon, Morrisville, NC, USA.

## Abstract

Mammalian embryos transiently exhibit aerobic glycolysis (Warburg effect), a metabolic adaptation also observed in cancer cells. The role of this particular type of metabolism during vertebrate organogenesis is currently unknown. Here, we provide evidence for spatio-temporal regulation of aerobic glycolysis in the posterior region of mouse and chicken embryos. We show that a posterior glycolytic gradient is established in response to graded transcription of glycolytic enzymes downstream of FGF signaling. We demonstrate that glycolysis controls posterior elongation of the embryonic axis by regulating cell motility in the presomitic mesoderm and by controlling specification of the paraxial mesoderm fate in the tail bud. Our results suggest that Warburg metabolism in the tail bud coordinates Wnt and FGF signaling to promote elongation of the embryonic axis.

## Introduction

Early studies in chicken and mouse embryos have established that energy metabolism is tightly regulated during development (Johnson et al., 2003; Spratt, 1948). The early mouse preimplantation embryo does not rely on glucose as its main source of energy but rather uses pyruvate and lactate to feed the Tricarboxylic Acid (TCA) cycle and produce ATP (Brinster, 1965). Around the time of implantation, a major metabolic transition occurs, leading the embryo to increase glucose uptake and glycolytic activity (Clough and Whittingham, 1983; Shepard et al., 1997). Most of this glycolytic activity coexists with an active TCA cycle and oxydative phosphorylation and results in lactate production (Johnson et al., 2003) thus resembling the Warburg metabolism or aerobic glycolysis of cancer cells (Vander Heiden et al., 2009). Subsequently, this intense glycolytic activity of the embryo decreases during organogenesis while respiration becomes the major mode of energy production (Wales et al., 1995). Aerobic glycolysis has been proposed to play a role in sustaining the intense proliferative activity of cancer and embryonic cells (Papaconstantinou, 1967; Vander Heiden et al., 2009). In the mammalian embryo, however, down-regulation of the glycolytic activity occurs during early organogenesis, when very high levels of proliferation are observed, thus questioning the role of this metabolic adaptation. To date, most studies of the metabolic status of vertebrate embryonic cells *in vivo* are based on metabolic tracing which can only provide crude spatial resolution. Recent studies have however shown that aerobic glycolysis can be regulated in a cell type and stage-specific manner as for instance in the developing retina, in osteoblasts or in endothelial cells (Agathocleous et al., 2012; Esen et al., 2013; Moussaieff et al., 2015). This raises the possibility of an instructive role for this particular type of metabolism in development (Shyh-Chang et al., 2013).

Here we investigated the role and regulation of metabolism in patterning and morphogenesis using musculo-skeletal development as a paradigm. Skeletal muscles and vertebrae derive from the paraxial mesoderm, which is continuously produced by gastrulation first in the primitive streak and then in the tail bud. Newly generated paraxial mesoderm appears as bilateral strips of mesenchyme called presomitic mesoderm (PSM) which periodically segment to generate the embryonic somites (Hubaud and Pourquie, 2014). Somites provide the blueprint for the metameric arrangement of vertebrae and associated muscles. Periodic somite formation is driven by a molecular oscillator, termed Segmentation Clock, which drives rhythmic activation of the Wnt, FGF and Notch pathways in the PSM (Hubaud and Pourquie, 2014). The segmental response to the oscillator is gated to a specific level of PSM called determination front by a system of traveling posterior to anterior gradients of Wnt and FGF signaling. Cells of the PSM exhibit a gradient of random motility (cell diffusion) controled by FGF, which has been proposed to control the posterior elongation movements involved in body axis formation (Benazeraf et al., 2010). In the paraxial mesoderm, the differentiation process is associated with a striking modular compartmentalization of the transcription of essential components of translation and oxidative metabolism which become upregulated as cells differentiate (Ozbudak et al., 2010). Furthermore, hypoxia can downregulate FGF signaling in the paraxial mesoderm leading to an arrest of the segmentation clock and ultimately to vertebral defects (Sparrow et al., 2012). These observations support a cross-talk between signaling and metabolism and argue for a dynamic regulation of metabolism during paraxial mesoderm development.

In this report, combining metabolomic and transcriptomic approaches, we identify a posterior to anterior gradient of glucose uptake and of Warburg type of metabolism in the tail bud region of the mouse and chicken embryos. We show that the tail bud glycolytic gradient is established in response to graded transcription of rate-limiting glycolytic enzymes downstream of FGF signaling. Inhibiting glycolysis in the chicken embryo leads to an arrest of axis elongation associated to an increased extracellular pH and decreased cell motility in the posterior PSM. The elongation arrest is also accompanied by premature differentiation of the SOX2-BRACHYURY neuro-mesodermal precursors (NMPs) toward a neural fate, resulting from inhibition of Wnt signaling in the tail bud. Thus our work identifies a striking role for glycolysis in integrating cell signaling during body axis formation.

## Results and discussion

### Identification of a posterior gradient of aerobic glycolysis in the mouse tail bud

To explore the regulation of metabolism during paraxial mesoderm differentiation, we performed a comprehensive metabolomic analysis of the developing posterior body axis in mouse embryos. To that end, we dissected the posterior part of 300 day 9.5 mouse embryos into 3 adjacent domains corresponding to the progressively more differentiated levels of the posterior PSM (P-PSM) including the tailbud, anterior PSM (A-PSM) and newly formed somites (Figure 1A). The relative abundance of a set of 2400 metabolites was analyzed for each sample by tandem mass spectrometry (LC-MS/MS and GC-MS). This strategy identified a total of 129 metabolites which were reliably detected in the embryo samples (Supplemental Figure 1). Thirty nine of these metabolites were differentially regulated during differentiation (Figure 1B, supplemental Table 1). Several metabolites involved in glycolysis, such as lactate or Glucose 6 phosphate (G6P) were detected at significantly higher levels (1.35 and 1.49 fold respectively, p<0.05) in P-PSM compared to A-PSM (Figure 1B-D, Supplemental Table 1). Other glycolytic metabolites including Glucose, Fructose-6-phosphate (F6P), Glyceraldehyde 3-phosphate (G3P), and 3-Phosphoglycerate (3-PG) also exhibited trends suggesting enrichment in the P-PSM (Figure 1 C-D, Supplemental Table 1). Other important nutrients such as Glutamine also showed a similar posterior gradient (Figure 1B). Using an enzymatic assay, we found significantly higher lactate levels in the P-PSM and tail bud, compared to A-PSM and somites (Figure 1E), consistent with more active glycolysis in more posterior regions. We also detected Cytochrome C oxidase activity in these posterior regions experiencing high glycolytic activity (Figure 1F). Activity of this enzyme which catalyzes the last step of the electron transfer chain requires oxygen, suggesting that lactate production occurs in aerobic conditions. Accordingly, exposing developing mouse embryos to hypoxic conditions disrupts FGF signaling and segmentation(Sparrow et al., 2012). Cytochrome C oxidase activity forms a gradient opposite to glycolysis, consistent with the increased regulation of translation and oxidative metabolism reported in the zebrafish anterior PSM (Ozbudak et al., 2010). Importantly, we did not detect any significant difference in ATP amount along the PSM, suggesting that the spatio-temporal regulation of glycolysis does not directly impact energy production (Figure 1G). We next analyzed the expression of key glycolytic enzymes in a previously generated PSM microarray series of consecutive micro-dissected fragments spanning the entire PSM in day 9.5 mouse embryos (Chal et al., 2015). Transcripts coding for *(Phosphoglucomutase (Pgm1/2), Hexokinase (HK1), Aldolase (AldoA), Triose phosphate isomerase (Tpi1),2,3 Phosphoglycerate mutase (Bpgm), Enolase (Eno3), Lactate dehydrogenase (Ldhb*) are enriched at the level of the tail bud/posterior-most PSM and show a posterior expression gradient (Figure 1H, Supplemental Table 2). Glucose transporter 3 (Glut3 or Slc2a3), which controls entry of glucose in the cell, also shows graded expression in the PSM and the tail bud at the mRNA and at the protein level (Figure 1H-I, Supplemental Table 2). Together, these observations identify a gradient of aerobic glycolytic activity associated to graded transcription of glycolytic enzymes in the posterior PSM and tail bud of mouse embryos.

**Figure 1:**
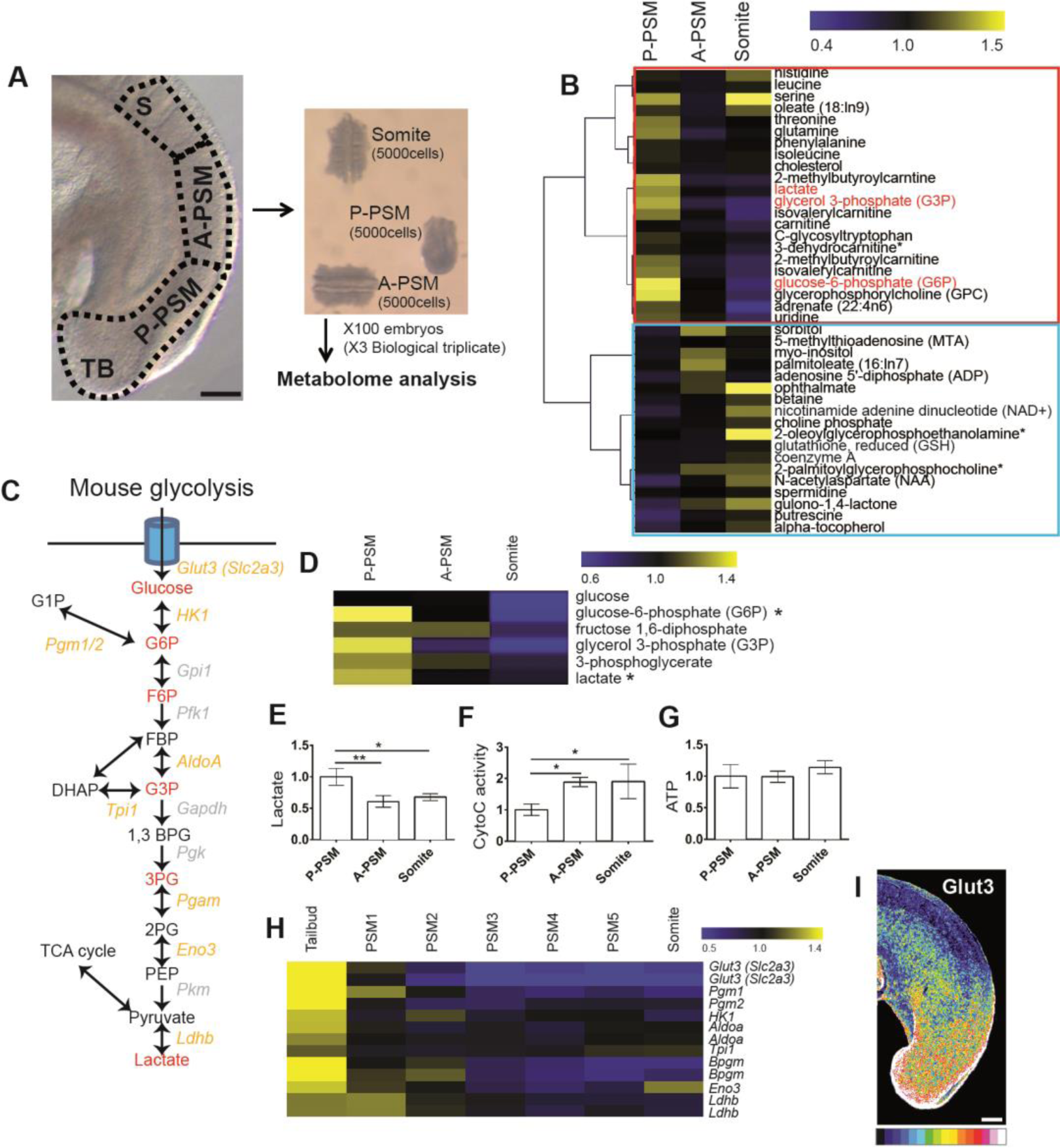
A posterior glycolytic gradient in the mouse tail bud. A-Lateral view of the posterior region of a 9.5 day mouse embryo showing the 3 domains used for metabolomics analysis. TB: Tail bud, S: somite, A-PSM and P-PSM: Anterior and Posterior Presomitic Mesoderm respectively. Each domain contains approximately 5000 cells. Scale bar: 100 μm. B-Clustering analysis of the 39 metabolites showing the most significant differential expression along the posterior region of the embryo. Color bar shows fold change from mean. Red box highlights metabolites downregulated and blue box metabolites upregulated during differentiation. C-Schematic representation of the mouse glycolytic cascade. Red labeling indicates metabolites detected in the metabolomics analysis. Yellow labeling indicates the glycolytic enzymes downregulated in the mouse embryo tail bud and PSMs. D-Expression profiles of the metabolites associated to glycolysis detected in the metabolomics analysis (* Significant at p<0.05 with t-test). The color bar shows fold change from mean of all triplicate samples. (E-G) Enzymatic detection of relative lactate levels, Cytochrome C oxidase activity, and relative ATP levels during tail bud differentiation. Graphs show triplicate experiments. Values were normalized by P-PSM, Error bars are ±SD. Statistical significance was assessed with one way ANOVA followed by Tukey’s test, * p<0.05, ** p<0.01. H- Expression profiles of transcripts coding for glycolytic enzymes downregulated in the mouse PSM during differentiation. Each value is normalized to the mean of MAS values of all triplicate mouse microarray series. The color bar shows fold change from mean. I-Lateral view of the tail bud region of 9.5-day mouse embryo stained with an antibody against Glut3 (n=5). Maximum projection of confocal sections. Fluorescence intensity is shown by pseudo-color image (16 color) using image J. Higher levels of Glut3 proteins are indicated in red-yellow. Scale bar: 100 μm.

### Conservation of the posterior glycolytic gradient in developing chicken embryos

We next investigated whether this graded glycolytic activity is conserved between mouse and chicken embryos. We generated a microarray series from consecutive fragments of the developing chicken PSM similar to that performed in mouse (Chal et al., 2015)(Figure 2A, Supplemental Figure 2). Most genes coding for glycolytic enzymes showed a posterior expression gradient in the chicken microarray series (Figure 2A-C, Supplemental Table 3). Graded expression of several glycolytic enzymes was confirmed by *in situ* hybridization (Figure 2D). The glucose transporter *GLUT3* was not detected in the chicken PSM but *GLUT1* (*SLC2A1*) shows a posterior expression gradient (Figure 2B-D, Supplemental Table 3). Furthermore, glucose uptake analysis with fluorescent glucose, (2NDBG) (Itoh et al., 2004; Yoshioka et al., 1996) demonstrates a clear posterior gradient peaking in the tail bud (Figure 2D). In the chicken embryo, analysis of lactate production using an enzymatic assay also shows a posterior gradient (Figure 2E). As observed in mouse, cytochrome C oxidase activity is also detected in the posterior glycolytic regions suggestive of an aerobic status of these regions in the chicken embryo (Figure 2F). No significant difference in ATP levels was observed along the antero-posterior axis (Figure 2G). Thus, the posterior to anterior gradient of aerobic glycolytic activity in the tail bud is conserved between mouse and chicken embryos.

**Figure 2:**
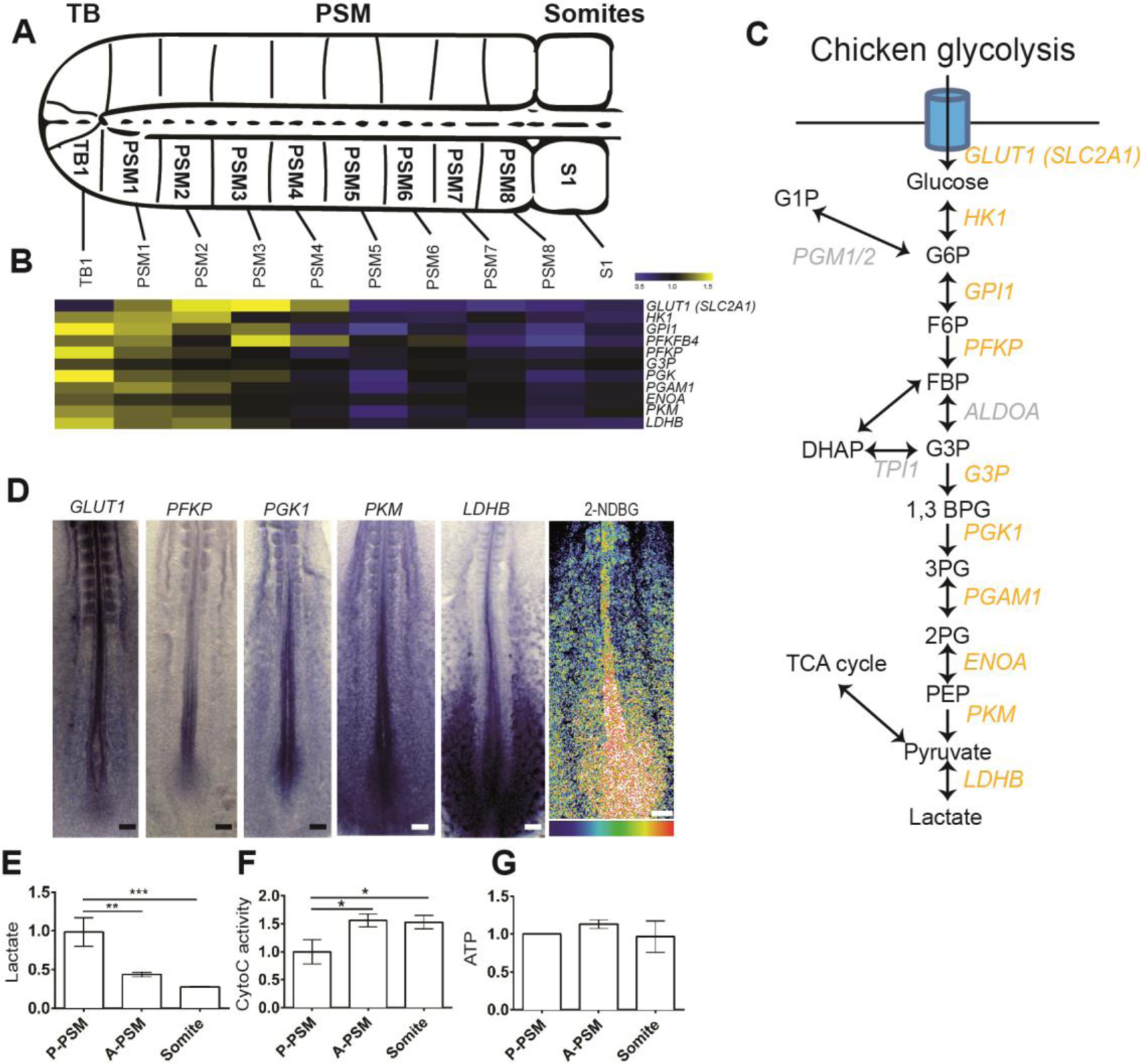
Conservation of the posterior glycolytic gradient in the chicken embryo. A- (Top). Schematic representation of the posterior portion of a two-day old chicken embryo. The fragments micro-dissected to generate the microarray series are indicated. Dorsal view, anterior to the right. TB: Tail bud, PSM: Presomitic Mesoderm. (Bottom)- B-Expression profiles of transcripts coding for glycolytic enzymes down-regulated in chicken PSM during differentiation as detected in the microarray series. Each value is normalized to the mean of MAS values of all duplicate chicken microarray series. Color bar shows fold change from mean. C-Schematic representation of the chicken glycolytic cascade. Yellow labeling indicates the glycolytic enzymes downregulated in the chicken embryo tail bud and PSMs. D-Left, posterior region of 2-day chicken embryos hybridized with probes for the glucose transporter *GLUT1*, and for the glycolytic enzymes *PFKP*, *PGK1*, *PKM* and *LDHB*. Right, fluorescent glucose (2-NDBG) uptake in the posterior region of a 2-day chicken embryo (n=11). Maximum projection of confocal sections. Fluorescence intensity is shown by pseudocolor image (16 color) using image J. Higher levels of fluorescent 2-NDBG are indicated in red-yellow. Ventral view, anterior to the top. Scale bar: 100 μm. (E-G) Enzymatic detection of relative lactate levels, Cytochrome C oxidase activity, and relative ATP levels during tail bud differentiation. Graphs represent triplicate experiments. Values were normalized by the P-PSM, Error bars ±SD. Statistical significance was assessed with one way ANOVA followed by Tukey’s test, * p<0.05, ** p<0.01, *** p<0.001).

### FGF signaling regulates the transcription of rate limiting glycolytic enzymes

High levels of aerobic glycolysis have often been associated with the need to produce important quantities of substrates for anabolic reactions required to sustain the rapid proliferation of cancer or embryonic cells (Vander Heiden et al., 2009). The proliferation rate and cell cycle length in the tail bud and in the somitic region have been measured using a variety of approaches and they were found to remain relatively stable in the 2-day chicken embryo with a duration of around 9-11 hours (Gomez et al., 2008; Primmett et al., 1989; Venters et al., 2008). Therefore, the spatio-temporal regulation of glycolysis in the trunk is unlikely to reflect changes in the proliferation regime of embryonic cells. In the 2-day old chicken embryo, fluorescent glucose uptake is strikingly regionalized, peaking in the tail bud, posterior PSM and neural tube, the forming limb buds, the rhombomere 4 region and the anterior neural ridge which are regions which largely overlap with regions where *FGF8* and its target *SPRY2* are expressed (Figure 3A-C). In the posterior region of the embryo, the gradient of glucose uptake and of glycolytic activity is parallel to the gradient of FGF signaling which controls segmentation and elongation of the body axis (Benazeraf and Pourquie, 2013). To investigate the interactions between FGF signaling and the regulation of glycolysis, we examined the metabolic activity following treatment with FGF/MAPK inhibitors (Figure 3D-F). Lactate production was inhibited following treatment with SU5402 (an FGFR1 inhibitor) and PD0359021 (a MAPK inhibitor) (Figure 3D). In contrast, these inhibitors had no effect on mitochondrial respiration or ATP production (Figure 3E-F). Inhibitors of other signaling pathways important for PSM patterning including Notch (DAPT) and Retinoic Acid (BMS204493) did not affect lactate or ATP production or cytochrome C oxidase function (Figure 3D-F). Significant down-regulation of the expression of genes coding for the rate-limiting glycolytic enzymes *PFKP*, *PGK1*, *PKM* and *LDHB* (0.83, 0.77, 0.73 and 0.52 fold respectively, p<0.01) was observed by qPCR after PD0359021 treatment compared to control untreated embryos (Figure 3G). Down-regulation of this subset of glycolytic enzymes after PD0359021 treatment was subsequently confirmed by *in situ* hybridization (Figure 3H-K, and not shown). These results suggest that the posterior gradient of FGF/MAPK signaling controls the high glycolytic activity in the posterior PSM/tail bud by regulating the transcription of rate-limiting glycolytic enzymes.

**Figure 3:**
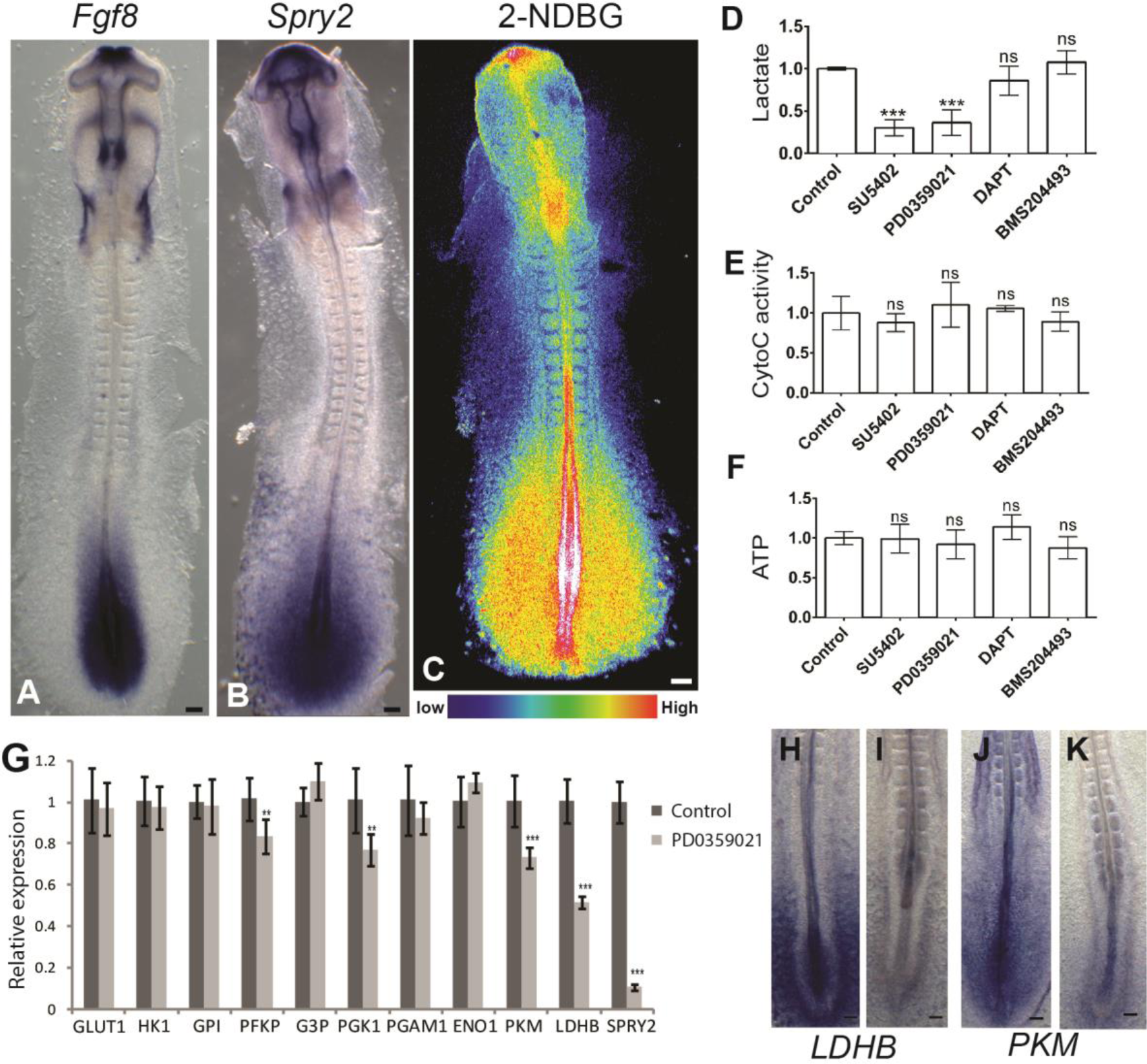
FGF signaling regulates glycolysis in the posterior part of the embryo. A-Fluorescent glucose (2NDBG) uptake in a 2-day old chicken embryo. Maximum projection of confocal sections. Fluorescence intensity is shown by pseudo-color image (16 color) using image J. Higher levels of uptake indicated in red-yellow. Dorsal view, anterior to the top. Scale bar: 100 μm. (B-C) Whole mount *in situ* hybridization of 2-day chicken embryos with *Fgf8* (B) and *Spry2*(C) probes. Dorsal view, anterior to the top. Scale bar: 100 μm. (D-F) Enzymatic detection of relative lactate levels (D), of Cytochrome C oxidase activity (E), and of relative ATP levels (F) in the posterior part of 2-day chicken embryos treated with inhibitors of FGF (SU5402), MAPK (PD0359021), Notch (DAPT) and Retinoic Acid (BMS204493) (n=6 for each condition). Graphs represent triplicate experiments. Values are normalized by untreated control embryos. Error bars ±SD. Statistical significance was assessed with one way ANOVA followed by Tukey’s test, ***p<0.001, ns p>0.05) G-qPCR analysis of glycolytic enzymes expression levels in control (blue) and in embryos treated with the MAPK-inhibitor PD0359021 (green). Graphs represent triplicate experiments. Values are normalized by untreated control embryos, Error bars are ±SD. Statistical significance was assessed with t-test, **p<0.01, ***p<0.001 (H-I) Whole mount *in situ* hybridization with probes for the rate limiting glycolytic enzymes *LDHB* (H: n=8, I: n=7) and *PKM* (N: n=8, O: n=8), in control (H, J) and PD0359021-treated (I, K) 2-day old chicken embryos. Ventral view, anterior to the top.

### Glycolysis regulates body axis elongation, extracellular pH and cell motility

To explore the role of glycolysis in tail bud development, we cultured stage 9-10 Hamburger and Hamilton (HH) (Hamburger and Hamilton, 1992) chicken embryos in the presence of the glycolysis inhibitor, 2-Deoxy-D-glucose (2DG), a competitive inhibitor of Hexokinase. 2DG treatment led to strong down-regulation of lactate production (Figure 4A), but not of cytochrome C oxidase activity in the embryo (Figure 4B). ATP production was not significantly changed upon 2DG treatment suggesting that glycolysis in the posterior PSM/tail bud plays a limited role in energy production (Figure 4C). In contrast, treating chicken embryos with the respiration inhibitor Sodium Azide (NaN_3_) did not affect lactate production while it effectively (albeit incompletely) inhibited Cytochrome C oxidase activity and ATP production in the tail bud region of the embryo (Figure 4A-C). 2DG treatment severely affected axis elongation, generating truncated embryos whereas somite formation continued at a normal pace leading to a progressive shortening of PSM length (Figure 4 D-F). In contrast, axis elongation was not affected by Sodium Azide treatment (Figure 4D, G), indicating a specific requirement of glycolysis for axial elongation.

**Figure 4:**
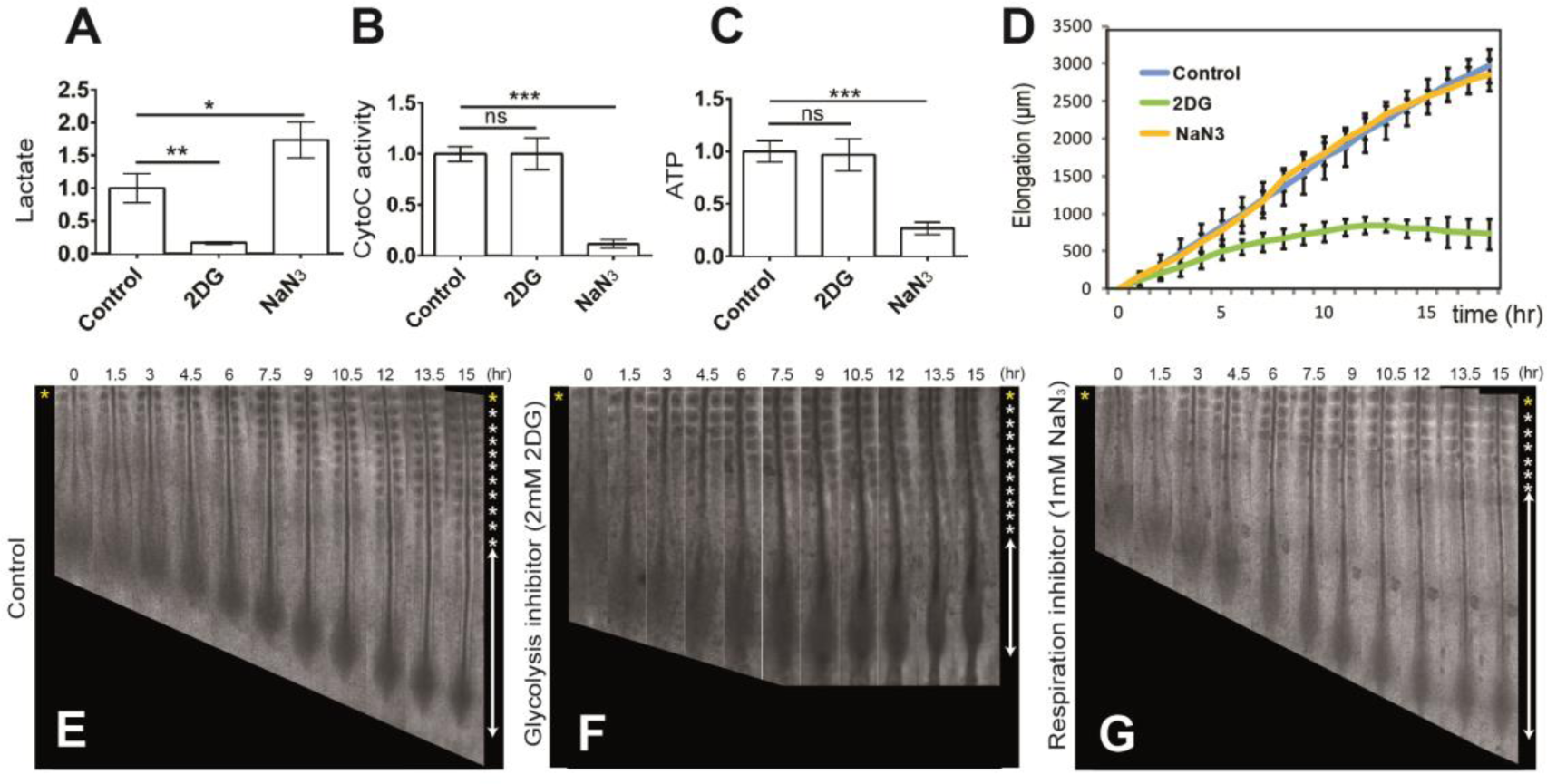
Glycolysis controls posterior elongation of the embryonic axis. (A-C) Enzymatic detection of relative lactate levels (A), of cytochrome C oxidase activity (B) and of relative ATP levels (C) in the posterior region of control 2-day old chicken embryos and in embryos treated with 2DG or NaN_3_ (n=6 embryos for each condition). Graphs represent triplicate experiments. Values are normalized by untreated control embryos. Error bars are ±SD. Statistical significance was assessed with one way ANOVA followed by Tukey’s test, **p<0.01, ***p<0.001, ns p>0.5) D- Increase in axis length (elongation) measured over time using time lapse microscopy (mean ±SD). Blue, control embryos (n=8); green, 2DG-treated embryos (n=7); yellow, NaN3-treated embryos (n= 5). (E-F) Elongation time course in a control (E), in a 2DG-treated (F) and in a NaN3-treated (G) 2-day chicken embryo. Bright field micrographs of the posterior region of a chicken embryos taken at 1.5 hour intervals. Somites formed at the last time point are indicated by asterisks on the right. Ventral views, anterior to the top. Scale bar: 500 μm.

A gradient of random cell motility (diffusion) controlled by FGF/MAPK signaling in the PSM has been proposed to drive posterior elongation movements in the chicken embryo (Benazeraf et al., 2010). To test whether glycolysis controls motility in the posterior PSM, we measured the diffusion and analyzed cell trajectories of posterior PSM cells labeled with H2B-GFP by electroporation in embryos treated or not with 2DG (Figure 5 A-C, and Supplemental movie 1). We show that inhibition of glycolysis leads to a progressive down-regulation of cell diffusion which correlates with the slowing down of elongation (Figure 5 D-F).

**Figure 5:**
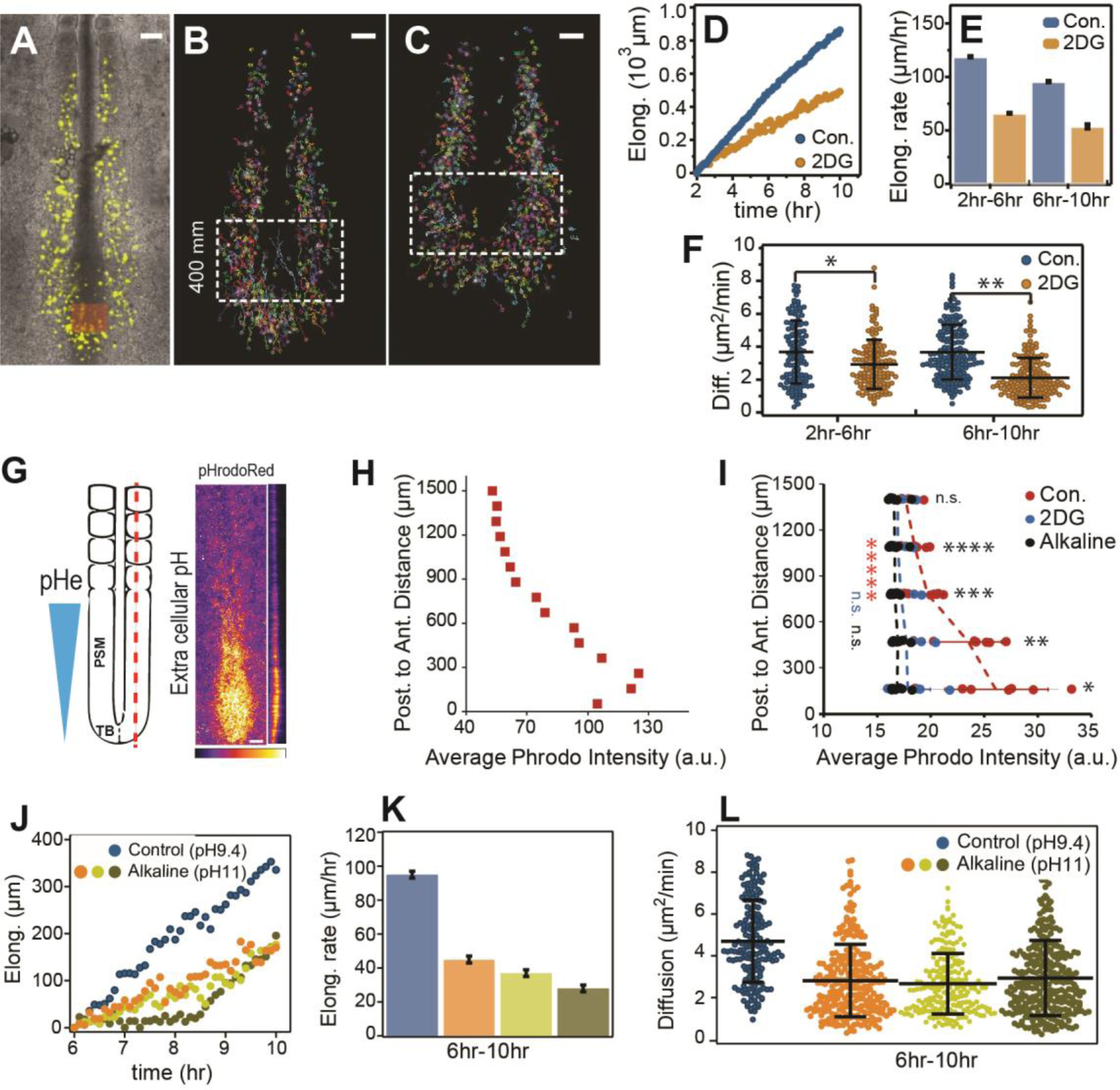
Glycolysis inhibition decreases cell motility and increases pH in the posterior PSM. (A-F) Effect of 2DG treatment on cell motility (diffusion) and PSM elongation. (A) Electroporated PSM cells expressing H2B-GFP are shown in yellow. (B) and (C) PSM cell trajectories for control and 2DG treated embryos respectively. Only tracks of cells located in the posterior PSM inside the box are used in the analysis. Scale bars: 100 μm. (D) Elongation curves showing the posterior displacement of the tail bud (orange box) as a function of time in the wild-type and 2DG-treated embryos. (E) Elongation rates of the embryos shown in (D), error bars are ±SD of the linear adjustments to data in (D). (F) Cell diffusion parameter for control and 2DG- treated embryos shown from 2h-6h and from 6h-10h (mean ±SD, *p=0.0004, **p< .0001, t-test). (G) - Extracellular pH in the posterior part of a 2-day old chicken embryo labeled with pHrodo Red showing the gradient of extracellular pH (pHe). Maximum projection(left) and sagittal section (right) of confocal sections of a 2-day chicken embryo labeled with pHrodo Red (right panel). Fluorescence intensity is shown by pseudocolor image (Fire color) using image J. Lower values of pH indicated in white-yellow. (H) - Quantification of intensity of pHrodo Red shown in (G). Each dot represents average fluorescence intensity measured in a ~0.06mm^2^ area of one embryo along the posterior to anterior axis. (I) - Effect of culturing embryos on 2DG-containing and on alkaline plates on the pHe gradient. Each dot represents average fluorescence intensity measured in a ~0.26mm^2^ area of each embryo along the posterior to anterior axis. Lines represent average intensity (±SD) (Control: n=7, alkaline: n=8, 2DG; n=7). (*p=0.001, **p=0.001, ***p=0.003, ****p=0.006, (n.s.) p=0.15, t-tests). ******(red, Con.) p=0.002, n.s.(blue, 2DG) p=0.14, n.s.(black, NaOH) p=0.35. Paired t-tests to compare the A-P ends of the pHe gradient. (J-L)- Effect of culture on alkaline plates on cell motility and PSM elongation. (J) Elongation curves for control and embryos cultured on alkaline plates. (K) Elongation rate corresponding to the curves in (J). (L) Cell diffusion for control and embryos cultured on alkaline plates (mean ±SD, p<.01 between control and NaOH cases and n.s. among NaOH using one way ANOVA followed by Tukey’s test).

In tumors or in blastocysts embryos, cell motility is promoted by aerobic glycolysis which leads to acidification of the extracellular environment as a result of lactic acid excretion (Gardner, 2015; Parks et al., 2013). This in turn which triggers activation of matrix metalloproteinases (MMP) involved in remodelling extracellular matrix. In the PSM, the pH sensitive enzyme MMP2 is expressed in a posterior gradient (Supplemental Figure 3). We observed that 2-day chicken embryos labeled with the pH sensor pHrodo Red (Ogawa et al., 2010) exhibit a posterior to anterior gradient of extracellular pH, with the lowest pH found in the tail bud where glycolysis is most active (Figure 5 G-H). This pH gradient can be abolished by incubating embryos on alkaline plates (pH 11) (Figure 5I). Treatment of the embryos with 2DG prior to labeling also results in a uniform higher pH (Figure 5I) suggesting that glycolysis is involved in the acidification of the extracellular environment at the posterior end of the embryo. Embryos cultured on alkaline plates also exhibit much slower axis elongation (Figure 5J) while somite formation continues to proceed at a normal pace (Supplemental Figure 3 and supplemental movie 2). Analysis of embryos in which paraxial mesoderm precursors were electroporated with an H2B-GFP construct cultured on alkaline plates revealed a decrease in cell diffusion paralleling the decrease in the speed of axis elongation (Figure 5J-L, supplemental movie 2). Together, these experiments suggest that aerobic glycolysis acts downstream of FGF signaling to promote acidification of the extracellular environment and cell motility.

### Glycolysis regulates Wnt signaling and maintenance of the NMPs in the tail bud

Rhythmic somite formation is driven by a molecular oscillator, termed segmentation clock, which drives cyclic activation of the Wnt, FGF and Notch pathways (Hubaud and Pourquie, 2014). Dynamic patterns of *Lunatic Fringe*, a cyclic gene controled by the segmentation clock were observed in 2DG-treated embryos suggesting that periodic signaling driven by the clock remains functional (Supplemental Figure 4). In the chicken embryo, the presumptive segment is first visible as a stripe of expression of the gene *CMESO1* (a chicken homologue of the mouse *Mesp2* gene). Upon 2DG treatment, *CMESO1* mRNA expression was normally expressed as a bilateral stripe in the anterior PSM (Figure 6 E, J). Thus inhibiting glycolysis impairs body elongation but not segmentation.

**Figure 6:**
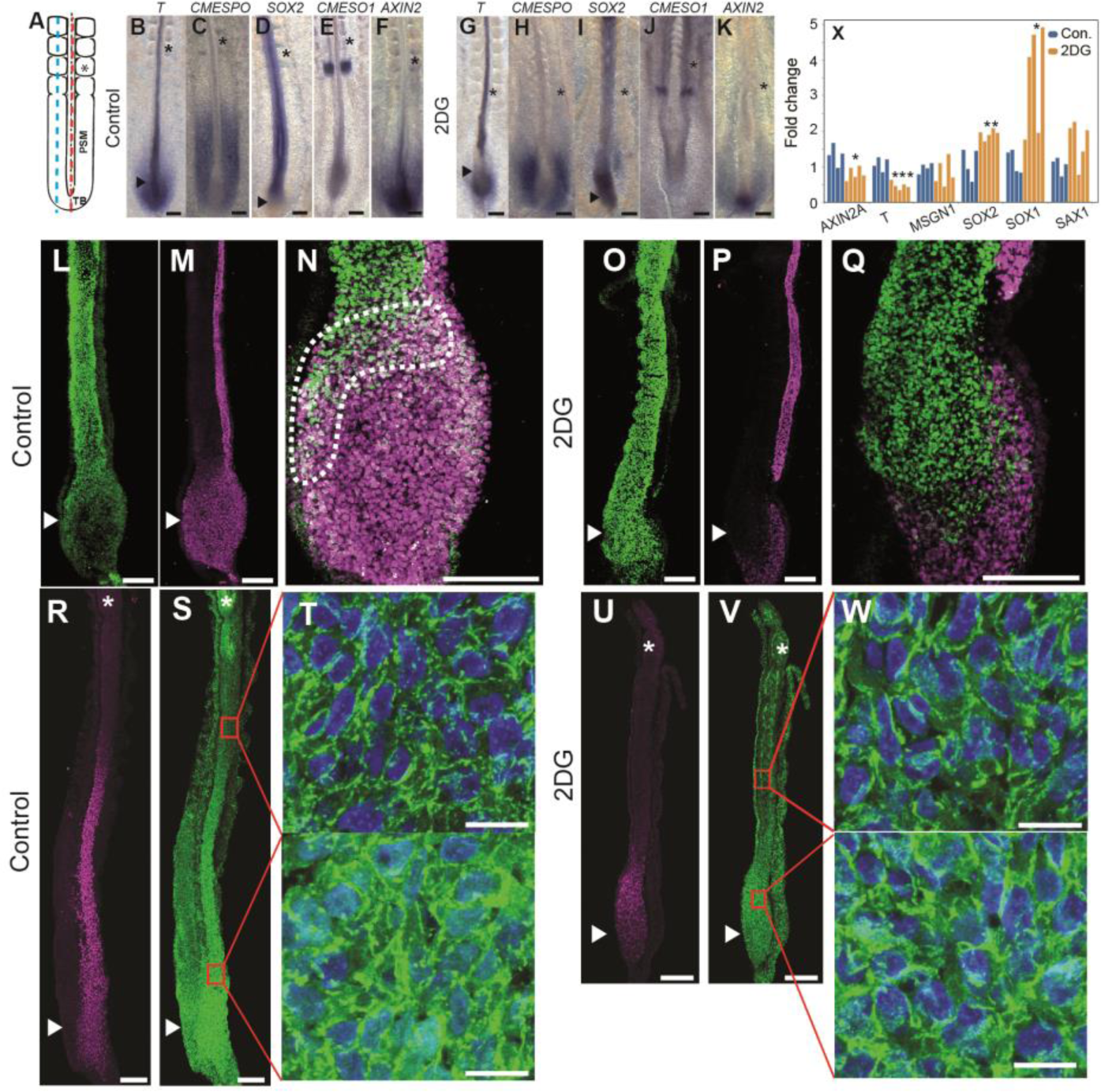
Inhibition of glycolysis phenocopies Wnt signaling inhibition in the tail bud. A-Schematic representation of the posterior part of a 2-day chicken embryo showing the position of the sections shown in L-Q (axial red dotted line) and R-W (Paraxial blue dotted line). (B-K) Whole mount *in situ* hybridizations showing the posterior region of 2-day old control (B-F) or 2DG-treated (G-K) chicken embryos hybridized with the following probes: *T*(*BRACHYURY*)(B: n=4, G: n=6); *CMESPO* (C: n=5, H: n=5); *SOX2* (D: n=7, I: n=4); *CMESO1* (E: n=5, J: n=4); *Axin2* (F: n=5, K: n=4). Asterisks indicate the last formed somite, and arrowheads indicate tail bud region. Ventral view, anterior to the top. Scale bar: 100 μm. (L-Q) SOX2 (L, O) and T /BRACHYURY (M, P) protein expression in a control (L-N) and a 2DG-treated (O-Q) embryo (n=4 for each condition). (N, Q) Higher magnification of the tail bud stained with SOX2 and T/BRACHYURY. Stippled line delineates the double positive cells in the chordo-neural hinge region. (R-T) Expression of CMESPO (R) and CTNNB1 (B-CATENIN) (S) proteins in sagittal sections of a 2-day old control chicken embryo (n=5). (T) Higher magnification of the regions boxed in red in (S) showing the nuclear localization of B-CATENIN. Top panel shows anterior PSM. Nuclei labeled with DAPI are shown in blue. (U-W) Expression of C-MESPO (U) and CTNNB1 (B-CATENIN) (V) proteins in sagittal sections of a 2-day old 2DG-treated chicken embryo (n=4). (W) Higher magnification of the regions boxed in red in (V) showing the nuclear localization of CTNNB1 (B-CATENIN). Top panel shows anterior PSM. Nuclei labeled with DAPI are shown in blue. (X)-qPCR analysis of the posterior region of 2-day old chicken embryos after 10 hr treated or not with 2DG. * p<0.05, **p<0.01, ***p<0.001. Scale bar: 100 μm (B-G, H-I, L-M), 10 μm (J-K, N-O) and 500 μm (P, Q). Asterisk indicates newly formed somites, and arrowheads indicate tail bud region.

Maintenance of the posterior elongation movements requires a constant supply of motile cells in the forming posterior PSM (Benazeraf and Pourquie, 2013). In 2DG-treated embryos, the size of the expression domains of the posterior PSM markers, *T/BRACHYURY* (Figure 6 A, B, G), and *CMESPO* (the chicken homologue of *Mesogeninl*) (Figure 6 C, H) strongly decreased, suggesting an arrest of PSM cells production. In contrast, expression of the neural marker *SOX2* was expanded posteriorly in the tail bud of treated embryos (Figure 6 D, I). No significant effect of 2DG treatment on proliferation and apoptosis in the paraxial mesoderm could be detected (Supplemental Figure 5). The posterior paraxial mesoderm and neural tube derive from a group of neuro-mesodermal precursors (NMPs) found in the tail bud region which co-express SOX2 and BRACHYURY (Henrique et al., 2015; Kimelman, 2016; Tzouanacou et al., 2009)(Figure 4L-N). In chicken embryos treated with 2DG, the SOX2-BRACHYURY double positive expression domain was replaced by cells only expressing SOX2 (compare Figure 6 L-N with O-Q). This suggests that NMPs differentiated into neural cells, leading to the termination of paraxial mesoderm production. We electroporated the NMP region in the anterior primitive streak with an H2B-RFP construct in transgenic chicken embryos expressing GFP ubiquitously and monitored the fate of the descendants of the electroporated NMPs. In control embryos, NMPs contribute to both neural and paraxial mesoderm lineages whereas in embryos treated with 2DG, NMPs stop producing paraxial mesoderm cells while continuing to populate the forming neural tube (Supplemental movie 3). Since axis elongation is largely driven by the posterior paraxial mesoderm (Benazeraf et al., 2010), arrest of its production can contribute to explain the arrest of elongation. Therefore, the high glycolytic state in the tail bud is also required for the maintenance of the NMPs cells and for the regulation of the balance of the differentiation rate between paraxial mesoderm and neural lineages. This phenotype resembles that observed when either *Wnt3a* or its targets *Brachyury* and *Tbx6* are mutated in mouse where ectopic neural tissue forms instead of paraxial mesoderm (Chapman and Papaioannou, 1998; Greco et al., 1996; Nowotschin et al., 2012; Takemoto et al., 2011; Yamaguchi et al., 1999). In the chicken embryo, the termination of axis elongation is also associated with a decrease in Wnt signaling and with the differentiation of the SOX2+/BRACHYURY+ cells into SOX2+/BRACHYURY- cells (Olivera-Martinez et al., 2012). Thus, the phenotype elicited by 2DG treatment phenocopies Wnt loss of function in the tail bud of mouse and chicken embryos. This led us to examine whether glycolysis inhibition down-regulates Wnt signaling. In control chicken embryos, nuclear β-catenin (CTNNB1) shows a posterior gradient in the PSM as reported for mouse embryos (Figure 6R-T) (Aulehla et al., 2008). This high nuclear β-catenin domain coincides with the expression domain of *CMESPO* (Figure 6 R), a direct target of Wnt signaling in the PSM (Buchberger et al., 2000; Wittler et al., 2007). 2DG treatment strongly reduced nuclear ß- catenin localization in the posterior PSM region (Figure 6U-W). By qPCR on 2DG treated embryos’ posterior domain, we detect a significant down-regulation of the Wnt targets *AXIN2* and *BRACHYURY* whereas the neural markers *SOX2* and *SOX1* were significantly up-regulated (Figure 6X). Together, these data argue that glycolysis regulates Wnt signaling in the posterior PSM/tail bud of the chicken embryo.

Wnt and FGF signaling are known to mutually regulate each other in the PSM (Aulehla and Pourquie, 2010; Naiche et al., 2011). Accordingly, embryos treated with the FGF/MAPK inhibitor PD0325901 also stopped elongating and showed reduced phosphorylated MAPK and Wnt activity (Figure 7A-I). Glycolysis inhibition also strongly reduced expression of markers of FGF activation such as phosphorylated MAPK and *SPROUTY2* (Figure 7F-G, M-N). Since *Wnt3a* in the PSM controls *Fgf8* expression in the mouse embryo (Aulehla et al., 2003), our data supports the existence of a closed regulatory loop linking FGF and Wnt signaling via glycolysis in the PSM. From these results we conclude that glycolysis downstream of FGF signaling is required to maintain the Wnt gradient in the posterior PSM. In turn, the Wnt gradient is required to maintain FGF activation.

**Figure 7:**
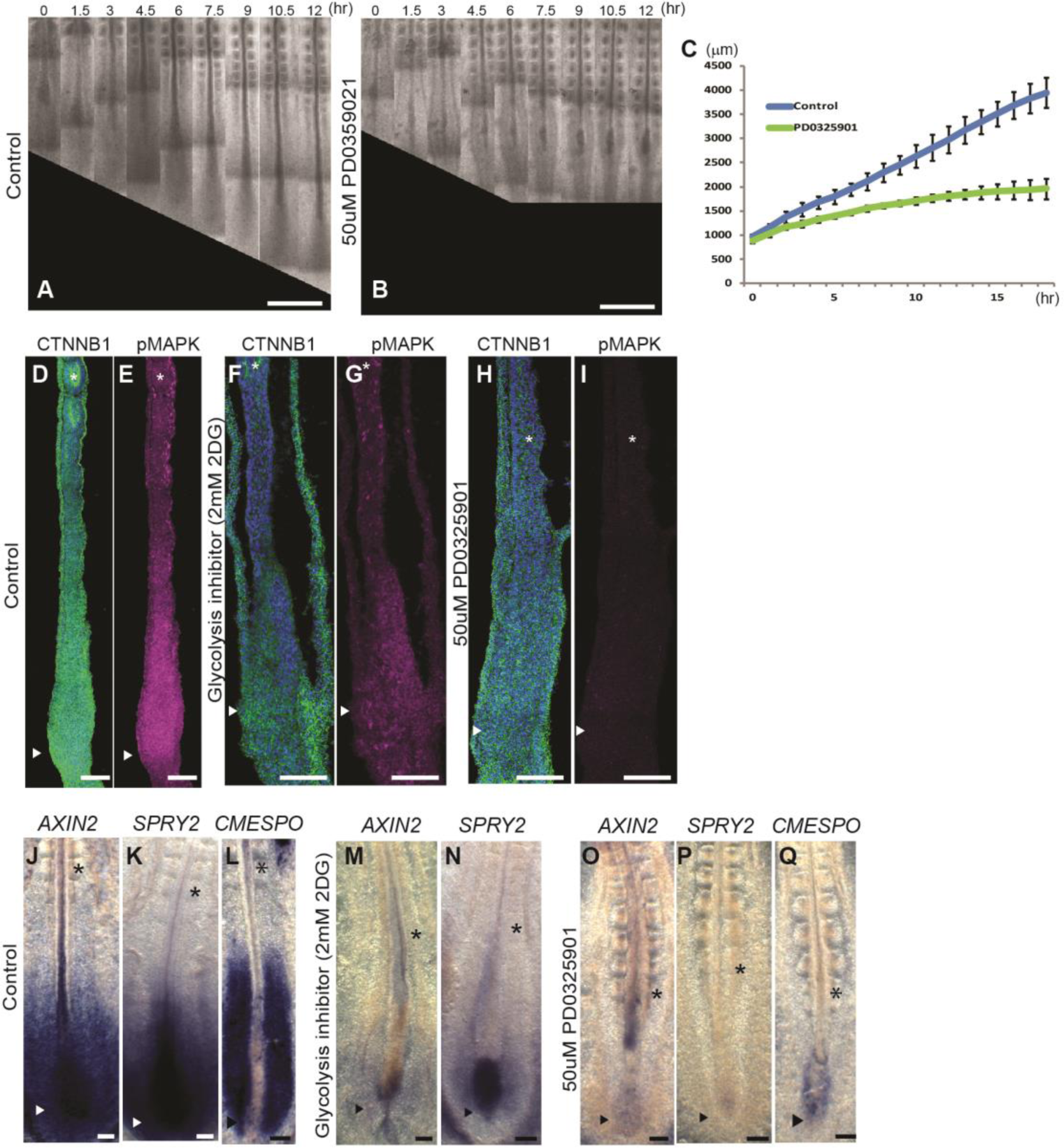
Phenotypic correlation between FGF and glycolysis inhibition. A-B Micrographs taken at 1.5 hour intervals of the posterior region of 2-day old chicken embryo control (A) and treated with PD0359021 (B). C- Graph showing the increase in axis length (elongation) measured using time lapse microscopy. in control embryos (blue) and PD0359021-treated embryo (green). Data represent the average of 5 embryos per conditions, error bars are mean ±SD. (D-I) Expression of CTNNB1 (Beta-CATENIN) (green, D, F, H) and phosphorylated MAPK (magenta, E, G, I), in longitudinal section of control (D, E: n=5), 2DG-treated (F, G: n=4) and PD0359021-treated (H, I: n=4) embryos. Sections shown in (D, F, H) are counterstained with DAPI (in blue) to visualize the nuclei. (J-Q) mRNA expression in the posterior part of 2-day old control of *AXIN2* (J, M, O), *SPRY2* (K, N, P), *CMESPO* (L, Q) detected by whole mount *in situ* hybridization (J: n=5, K: n=5, L: n=4), 2DG-treated embryo (M: n=4, N: n=3) and PD0359021-treated embryos (O: n=4, P: n=3, Q: n=4). Scale bar: 500 μm (A, B) 100 μm (D-Q). Ventral view, anterior to the top. Asterisk indicate newly formed somites, and arrowheads indicate tail bud region.

## Conclusion

Our study shows that glycolysis acts independently of energy production in specific developmental processes necessary to sustain posterior elongation of the embryonic axis: the Wnt-dependent production of paraxial mesoderm cells from tail bud NMPs and the control of their motility. In yeast and cancer cells, glycolysis regulates intracellular pH which in turn controls V-ATPase assembly (Dechant et al., 2010). The recent implication of V-ATPase in Wnt signaling might provide a link between glycolysis and Wnt in the embryo (Cruciat et al., 2010). The effect on cell motility might be linked to the role of localized glycolytic activity ensuring rapid delivery of ATP for actin polymerization to the forming protrusions which was hypothesized in cancer or in endothelial cells (De Bock et al., 2013; Nguyen et al., 2000). Interestingly, other developmental processes such as segmentation are not as sensitive to glycolysis inhibition. The striking similarities between this embryonic metabolic state and the Warburg metabolism suggest that cancer cells could redeploy a specific embryonic metabolic program with significant consequences on cell signaling and proliferation.

## Acknowledgements

We thank members of the Pourquié laboratory and Michel Labouesse, Norbert Perrimon, Alexander Aulehla and Cliff Tabin for discussions and comments on the manuscript. We are grateful to Bertrand Bénazéraf, Alexis Hubaud, Nicolas Denans, and Aurélie Krol for assistance with some chicken embryo experiments. We thank members of IGBMC imaging and microarray facility, and Jean Marie Garnier for construction of plasmids. This work was supported by an advanced grant of the European Research Council, and a grant of the Fondation pour la Recherche Médicale (SPF20120523860) to MO. F.X. was supported by a HHMI-HHWF fellowship. Microarrays data were deposited in GEO database under the accession number of GSE39613 for Mouse and GSE75798 for chicken.

## Authors’ contributions

M.O. designed, performed and analyzed the experiments with O.P. E.K. supervised the metabolomic analysis. P. M. analyzed the microarray data. J.C. generated the anti-CMESPO antibody. K.G. and F. X. analyzed the imaging data. M.O. and O.P. wrote the manuscript and O.P. supervised the project. All authors discussed and agreed on the results and commented on the manuscript.

## Experimental procedures

The posterior end of three hundred day 9.5 CD1 mouse embryos were dissected into three consecutive fragments. Metabolite content of pooled fragments of the same antero-posterior level was analyzed by tandem Mass Spectrometry as described in (Evans et al., 2009). Fertilized chicken eggs were obtained from commercial sources. Eggs were incubated at 38 °C in a humidified incubator, and embryos were staged according to Hamburger and Hamilton (HH) *(23)*. For the microarray series, both left and right posterior paraxial mesoderm from a stage 12HH chicken embryo were dissected in ten consecutive fragments which were each used to generate RNA hybridized to one Affymetrix microarray as described in (Chal et al., 2015). Analysis of the microarray series was performed as described in (Chal et al., 2015). In most experiments described in this study, we cultured chicken embryos at 38°C from Stage 9HH using the Early Chick (EC) culture system (Chapman et al., 2001). Albumin plates containing either 2mM 2DG (2-Deoxy-D-glucose; Sigma), 1mM NaN_3_ (sodium azide; Sigma), were prepared to respectively inhibit glycolysis and respiration. For signaling pathways inhibition, 500 μM SU5402(Sigma), 50 μM PD035901 (AXON MEDCHEM BV), 100 μM DAPT (Sigma) or 50 μM BMS204493 were diluted in PBS, and 100 μl of the inhibitor solution was added under and on top of the embryos. Cultured embryos were reincubated at 38C for various time periods and processed for *in situ* hybridization, immunohistochemistry or qPCR. In some cases, the paraxial mesoderm cells were electroporated before embryo culture as described in (Denans et al., 2015). Plasmids used for electroporation include H2B-RFP. Extracellular pH was measured by incubating embryos in the pHrodo Red pH Indicator (Lifetechnologies). 23mM NaOH albumin plates exhibiting a pH of 11.0-11.3 (the pH of control plates is 9.4-9.6) were used to test the effect of lowering the extracellular pH on pHrodo expression. The posterior elongation of the embryonic axis was measured using time lapse videomicroscopy. To quantify axis elongation, the last formed somite at the beginning of the time-lapse experiment was taken as a reference point (yellow asterisk in Fig. 3 E and F), and the position of the Hensen’s Node with respect to this somite was tracked as a function of time using the manual tracking plug-in in Image J. Enzymatic measurements of lactate (Biovision), ATP (Perkin Elmer), and cytochrome C activity (Sigma) were performed using commercial kits according to manufacturer’s instruction. Glucose uptake was measured by incubating embryos in 2-(*N*-(7-Nitrobenz-2-oxa-1,3-diazol-4-yl)Amino)-2-Deoxyglucose (2-NDBG, Lifetechnologies) followed by imaging on a confocal macroscope (Leica).

---

**Supplemental Figure 1, related to Figure 1.**
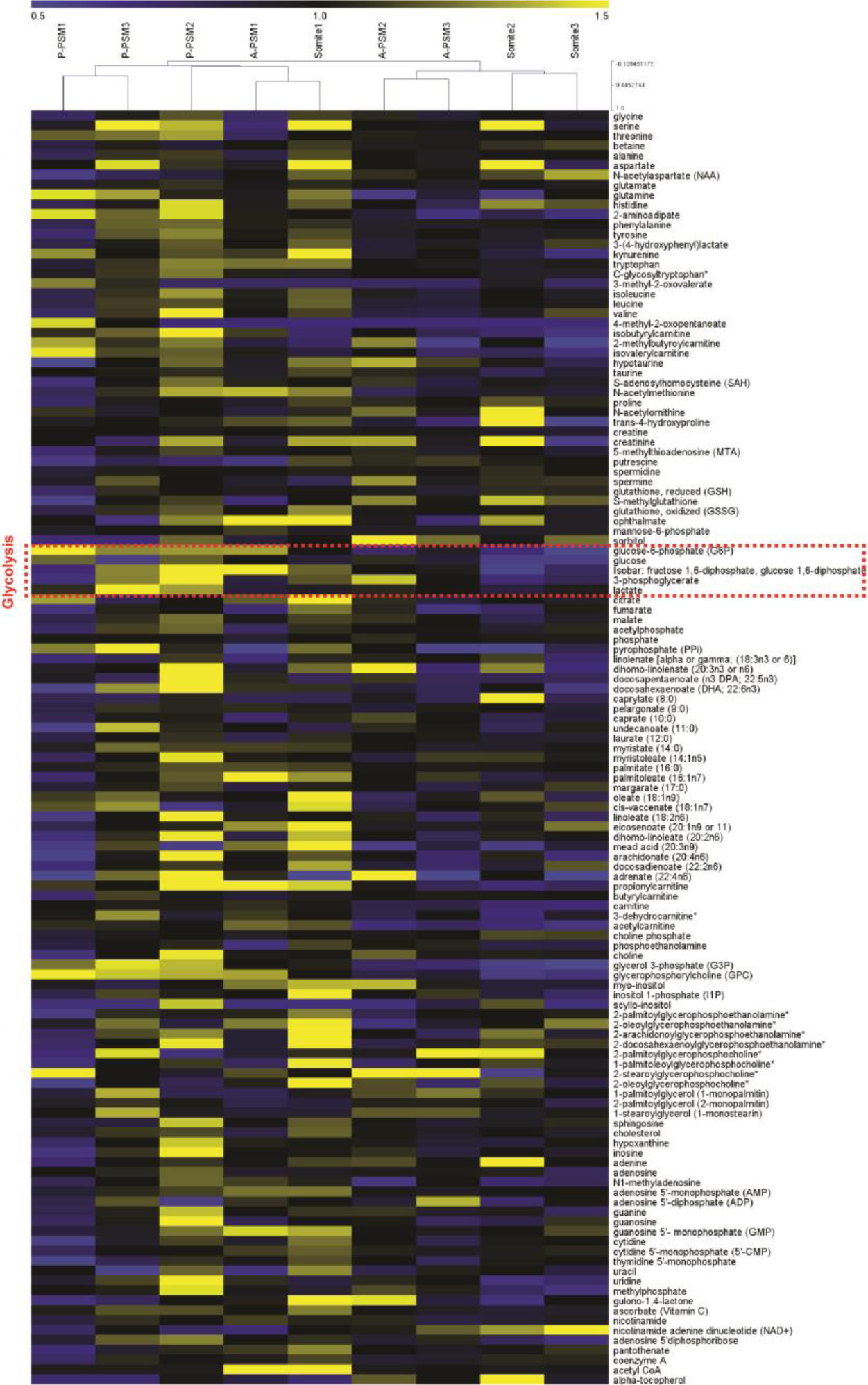
Hierarchical clustering of normalized mouse metabolome data. Color Heat map showing the fold change of the 129 metabolites identified by mass spectrometry analysis from biological triplicate samples of the 9.5 day mouse embryo posterior body region shown in Figure 1A. Each data point is normalized to the mean value of all triplicate samples. Hierarchical clustering of triplicate P-PSM, A-PSM samples, and 2 somite samples using Pearson correlation coefficient. Dotted red box highlights glycolysis metabolites.

**Supplemental Figure 2, related to Figure 2.**
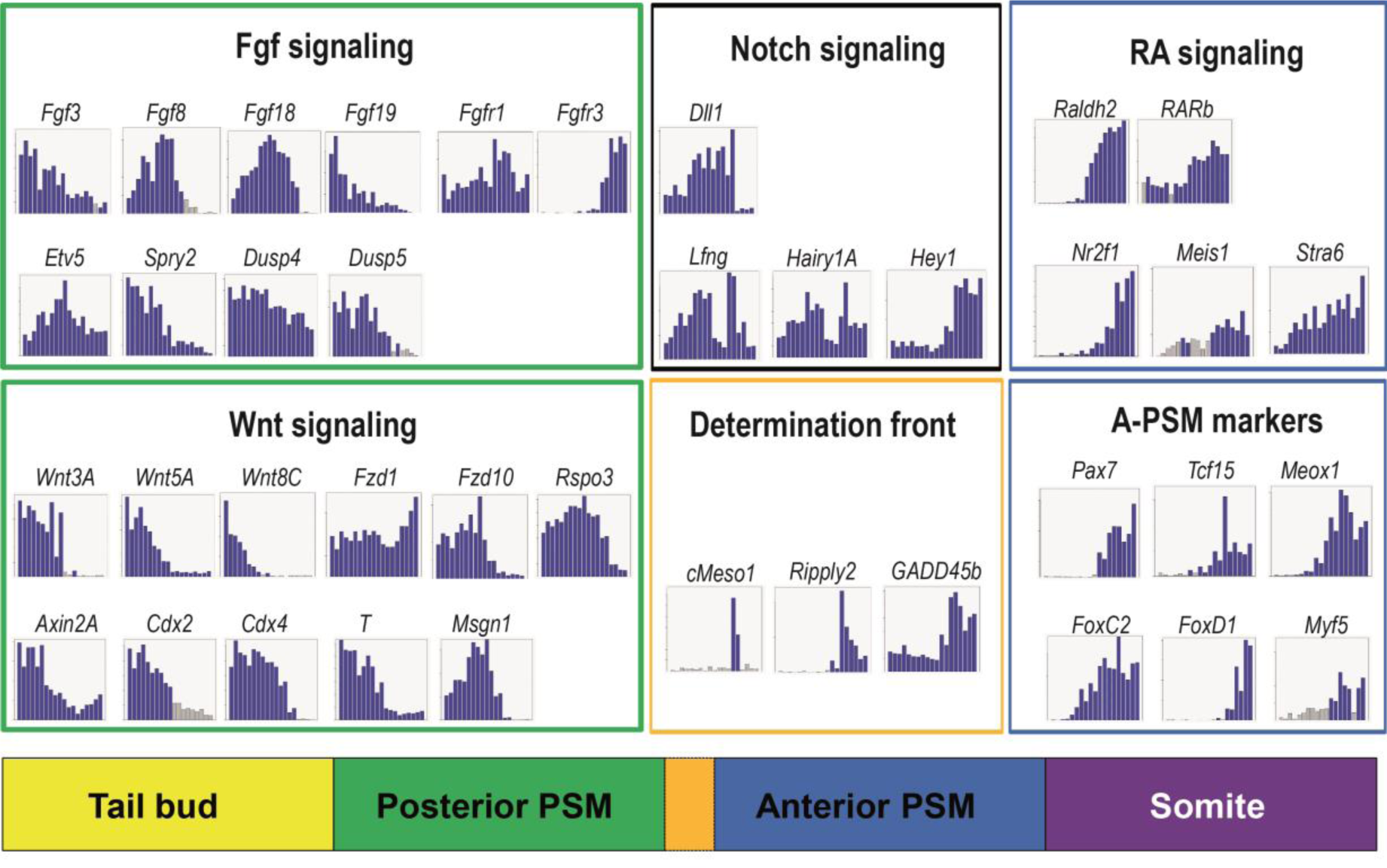
Generation of gene expression profiles in the chicken PSM differentiation. Histograms representing expression profiles of transcripts detected in the microarray series of chicken embryo posterior paraxial mesoderm described in Figure 2. The histograms were generated from MAS expression value for a given probeset, with samples of the two microdissection series arranged according to their position along the antero-posterior axis (posterior to the left) as described in Chal et al. (2015). The resulting profile gives a quantitative view of transcript expression during early stages of paraxial mesoderm differentiation. Expression of representative genes associated to major signaling pathways involved in PSM patterning and differentiation is shown. “Posterior” signaling pathway genes are highlighted by a Green box, and Blue box represent genes for anterior signaling pathways. Orange box show genes preferentially expressed at the determination front. Notch signaling pathway components including cyclic genes are highlighted by a Black box. Bottom: schematic representation of the signaling gradients and of the major posterior paraxial mesoderm domains (color-coded). Orange bar marks the determination front level where cells acquire their segmental identity.

**Supplemental Figure 3, related to Figure 5.**
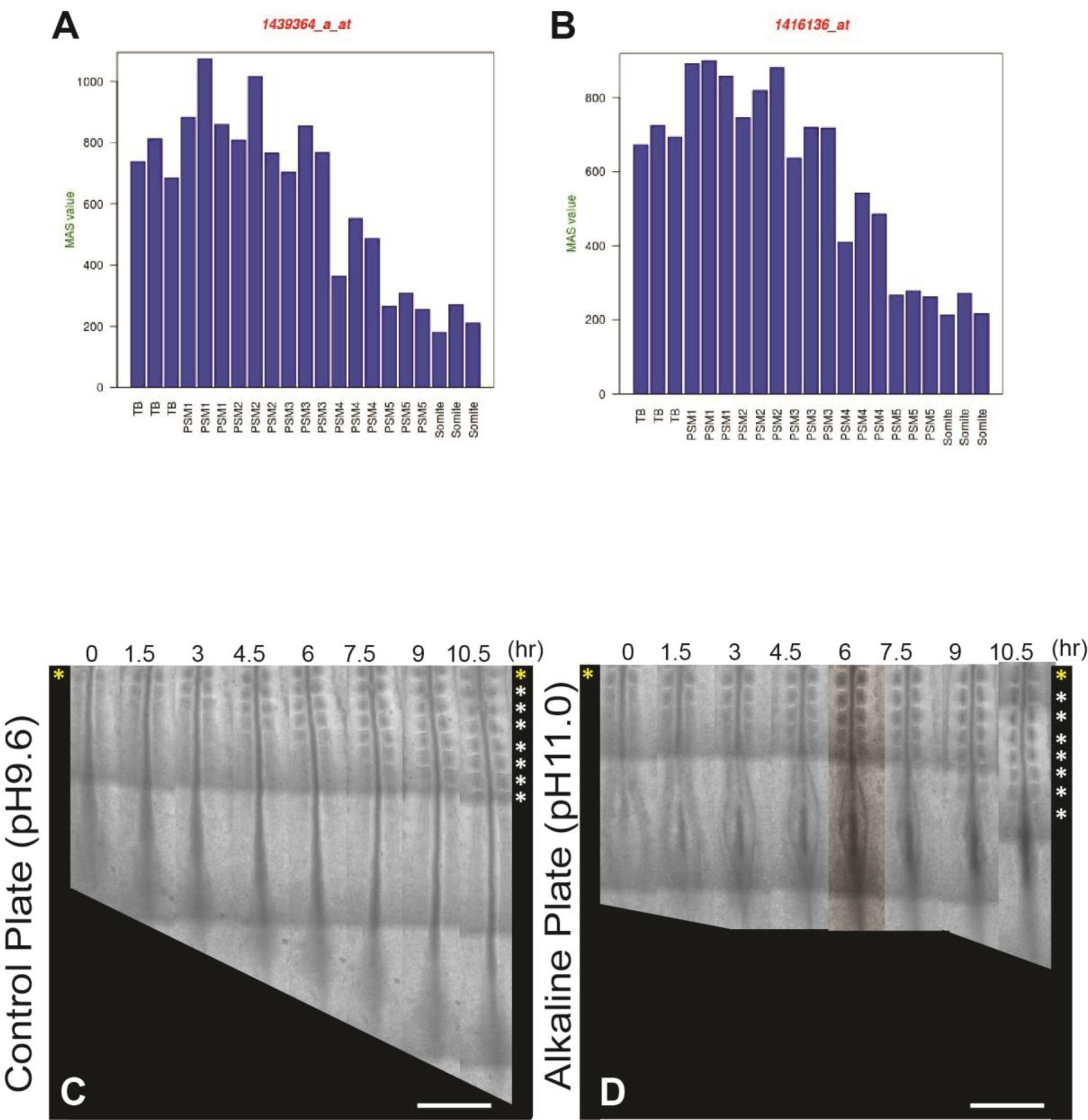
Effect of pH increase on elongation. (A, B) Expression dynamics of two different probesets of *MMP2* in the tail bud and posterior PSM as detected in the chicken microarray series for two different probesets. (C, D) Elongation time course in a control plate (C: pH 9.6) and in alkaline plate (D: pH11.0) chicken embryo. Bright field micrographs of the posterior region of a chicken embryos taken at 1.5 hour intervals. Somites formed at the last time point are indicated by asterisks on the right. Ventral views, anterior to the top. Scale bar: 500 μm.

**Supplemental Figure 4, related to Figure 6.**
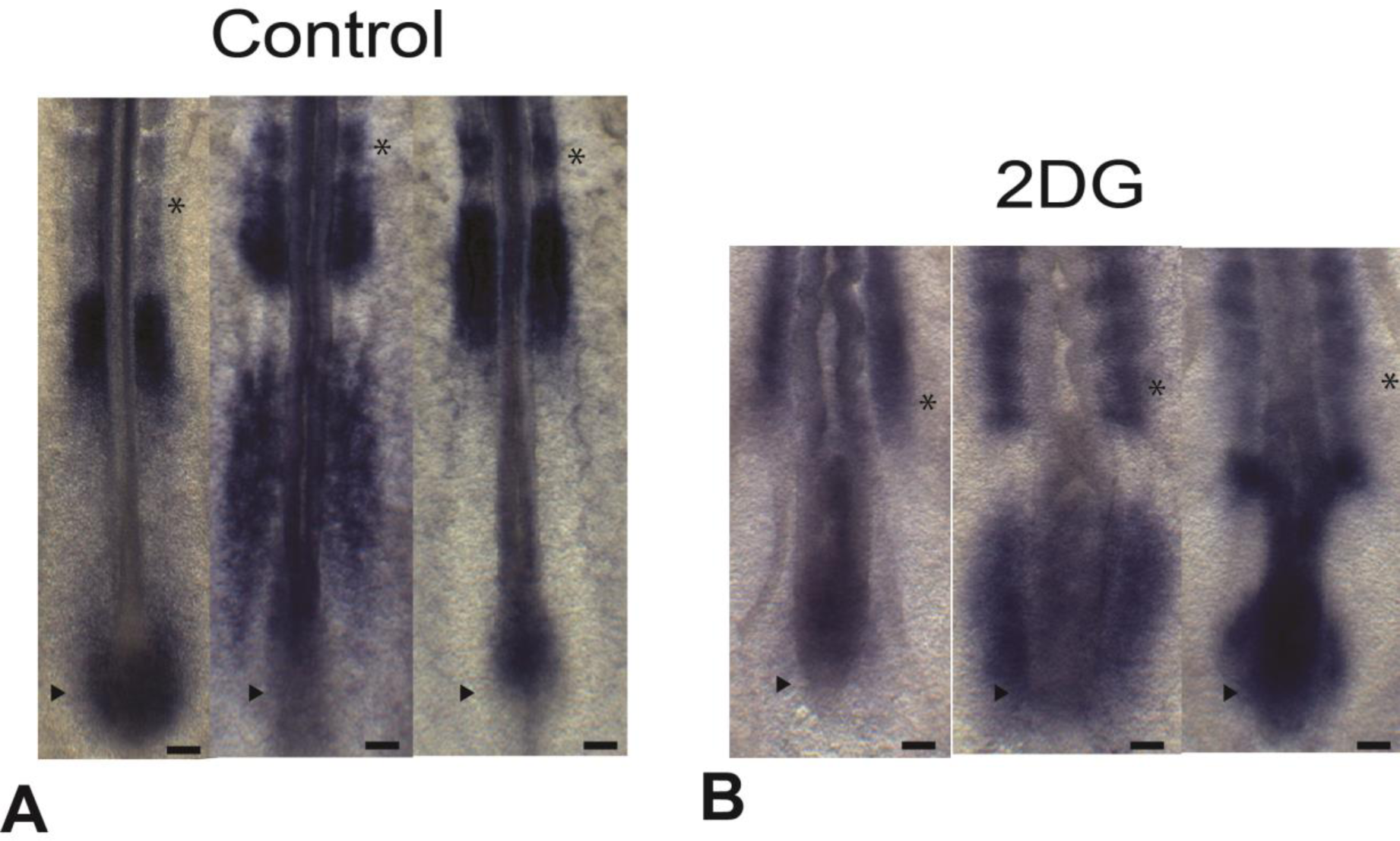
Glycolysis inhibition does not block cyclic gene expression. Whole mount *in situ* hybridizations showing the posterior region of 2-day old control (B-F) or 2DG-treated (G-K) chicken embryos hybridized with *L-fng* probe. Scale bar: 100 μm. Asterisk indicate newly formed somites, and arrowheads indicate tail bud region.

**Supplemental Figure 5, related to Figure 6.**
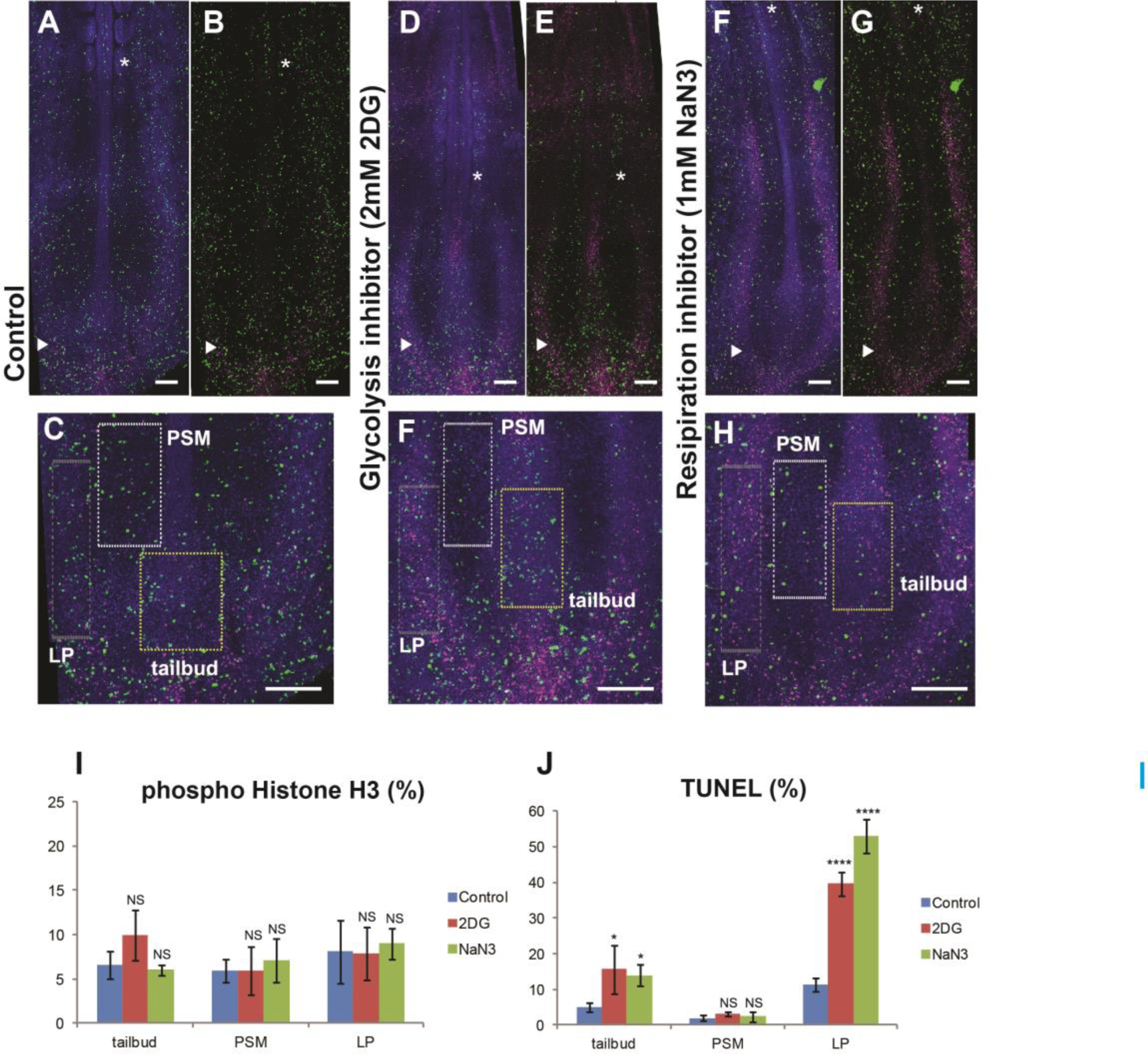
Axial elongation defects caused by glycolysis inhibition are independent of cell proliferation and apoptosis. (A-H) 2-day old chicken embryos stained in whole mount with phosphorylated histone H3 (pH3, green) for proliferating cells, and with TUNEL staining (magenta) for apoptotic cells. Nuclei are labeled with DAPI (blue). Control (A-C), 2DG (B-F), NaN3-treated embryo (FH). Ventral views, anterior to the top. Scale bar: 100 μm. Asterisk indicate newly formed somites, and arrowheads indicate tail bud region.(C, F, H)- Higher magnification of the panels shown in A-G showing the tail bud region. Dashed boxes show areas where proliferation and apoptosis were analyzed in the tailbud region (yellow line), the PSM region (white line) and the lateral plate mesoderm region (grey line). Scale bar: 100 μm.(I-J) Histograms showing the corresponding quantification of cell proliferation (I) and apoptosis (J) by mesoderm regions and by treatments. Data represent four independent experiments with error bars ± SD. Statistical significance was assessed with one way ANOVA followed by Tukey’s test (*P<0.05, ****P<0.0001, ns P>0.05).

**Supplemental Table 1, related to Figure 1.** Metabolomic analysis of the 9.5 day old mouse embryo tail bud and posterior region Fold change variations along the paraxial mesoderm of the 129 metabolites shown in Supplemental Figure 1. Pathways are assigned for each metabolite, also allowing examination of overrepresented pathways. Statistical comparisons are with both Student’s t-tests (p-value) and false discovery rate analysis (q-value). Significant metabolite amount differences in a given Paraxial mesoderm domain (*p* ≤ 0.05) are indicated by **bold** font values. Differences that not statistically significant but nevertheless close to the cutoff (0.05<p<0.10) are indicated in italic font values. Normal font values were found not significant.

| Metabolite |  |  | Fold of Change |  |  | Statistical Value |  |  |  |  |  |
| --- | --- | --- | --- | --- | --- | --- | --- | --- | --- | --- | --- |
| Name | Super Pathway | CAS | <u>P-PSM/</u><br><u>A-PSM</u> | <u>P-PSM/</u><br><u>somite</u> | <u>A-PSM/</u><br><u>somite</u> | P-PSM/<br>A-PSM |  | P-PSM/somite |  | A-PSM/somite |  |
|  |  |  |  |  |  | P-VAL<br>UE | Q-VAL<br>UE | P-VAL<br>UE | Q-VAL<br>UE | P-VAL<br>UE | Q-VAL<br>UE |
| glycine | Amino acid | 56-40-6; | 1.11 | 0.96 | 0.87 | 0.153 | 0.835 | 0.797 | 0.996 | 0.422 | 0.493 |
| serine | Amino acid | 56-45-1; | **1.49** | 0.87 | 0.58 | 0.046 | 0.591 | 0.817 | 0.996 | 0.254 | 0.391 |
| threonine | Amino acid | 72-19-5; | **1.37** | **1.27** | 0.92 | 0.049 | 0.591 | 0.024 | 0.565 | 0.474 | 0.509 |
| betaine | Amino acid | 107-43-7; | 0.93 | *0.82* | **0.88** | 0.418 | 0.994 | 0.069 | 0.810 | 0.013 | 0.223 |
| alanine | Amino acid | 56-41-7; | 1.08 | 0.95 | 0.88 | 0.423 | 0.994 | 0.746 | 0.996 | 0.376 | 0.486 |
| aspartate | Amino acid | 56-84-8; | 1.23 | 0.83 | 0.68 | 0.228 | 0.839 | 0.758 | 0.996 | 0.337 | 0.462 |
| N-acetylaspartate (NAA) | Amino acid | 997-55-7;997-55-7 ; | *0.72* | *0.59* | 0.83 | 0.086 | 0.617 | 0.056 | 0.798 | 0.124 | 0.315 |
| glutamate | Amino acid | 56-86-0; | 1.03 | 1.00 | 0.97 | 0.752 | 0.994 | 0.954 | 0.996 | 0.324 | 0.459 |
| glutamine | Amino acid | 56-85-9; | **1.59** | 1.35 | 0.85 | 0.012 | 0.591 | 0.117 | 0.933 | 0.218 | 0.389 |
| histidine | Amino acid | 5934-29-2; | 1.16 | 0.87 | **0.75** | 0.549 | 0.994 | 0.386 | 0.933 | 0.040 | 0.225 |
| 2-aminoadipate | Amino acid | 542-32-5;1118-90-7; | **1.61** | **1.67** | 1.03 | 0.031 | 0.591 | 0.022 | 0.565 | 0.488 | 0.513 |
| phenylalanine | Amino acid | 63-91-2; | 1.12 | 1.06 | **0.94** | 0.646 | 0.994 | 0.857 | 0.996 | 0.032 | 0.225 |
| tyrosine | Amino acid | 60-18-4; | 1.08 | 1.05 | 0.98 | 0.873 | 0.994 | 0.942 | 0.996 | 0.780 | 0.599 |
| 3-(4-hydroxyphenyl)lactate | Amino acid | 6482-98-0; | 1.08 | 0.94 | 0.87 | 0.725 | 0.994 | 0.756 | 0.996 | 0.123 | 0.315 |
| kynurenine | Amino acid | 2922-83-0; | 1.15 | 1.03 | 0.89 | 0.209 | 0.835 | 0.648 | 0.995 | 0.977 | 0.646 |
| tryptophan | Amino acid | 73-22-3; | 1.01 | 1.03 | 1.02 | 0.983 | 0.994 | 0.907 | 0.996 | 0.519 | 0.516 |
| C-glycosyltryptophan* | Amino acid |  | 1.09 | 1.20 | **1.10** | 0.525 | 0.994 | 0.206 | 0.933 | 0.001 | 0.085 |
| 3-methyl-2-oxovalerate | Amino acid | 51829-07-3; | 1.39 | 1.28 | 0.92 | 0.199 | 0.835 | 0.447 | 0.933 | 0.423 | 0.493 |
| isoleucine | Amino acid | 73-32-5; | 1.14 | 1.03 | *0.91* | 0.595 | 0.994 | 0.935 | 0.996 | 0.065 | 0.299 |
| leucine | Amino acid | 61-90-5; | 1.11 | 0.98 | **0.88** | 0.605 | 0.994 | 0.881 | 0.996 | 0.019 | 0.225 |
| valine | Amino acid | 72-18-4; | 1.27 | 1.05 | 0.83 | 0.539 | 0.994 | 0.975 | 0.997 | 0.146 | 0.331 |
| 4-methyl-2-oxopentanoate | Amino acid | 816-66-0; | 1.54 | 1.54 | 1.00 | 0.220 | 0.835 | 0.220 | 0.933 |  |  |
| isobutyrylcarnitine | Amino acid | 25518-49-4; | 1.30 | **1.84** | **1.41** | 0.218 | 0.835 | 0.027 | 0.565 | 0.034 | 0.225 |
| 2-methylbutyrylcarnitine | Amino acid | 31023-25-3; | 1.37 | *1.65* | *1.20* | 0.211 | 0.835 | 0.075 | 0.810 | 0.089 | 0.315 |
| isovalerylcarnitine | Amino acid |  | 1.37 | 1.69 | 1.23 | 0.052 | 0.591 | 0.006 | 0.408 | 0.032 | 0.225 |
| hypotaurine | Amino acid | 300-84-5; | 0.71 | 0.81 | 1.13 | 0.331 | 0.994 | 0.583 | 0.961 | 0.611 | 0.529 |
| taurine | Amino acid | 107-35-7; | 1.08 | 1.11 | 1.04 | 0.152 | 0.835 | 0.463 | 0.933 | 0.786 | 0.599 |
| S-adenosylhomocysteine (SAH) | Amino acid | 979-92-0; | 0.95 | 0.97 | 1.02 | 0.753 | 0.994 | 0.773 | 0.996 | 0.868 | 0.619 |
| N-acetylmethionine | Amino acid | 65-82-7; | 1.06 | 1.04 | 0.99 | 0.871 | 0.994 | 0.921 | 0.996 | 0.716 | 0.584 |
| proline | Amino acid | 147-85-3; | 0.90 | 0.79 | 0.88 | 0.174 | 0.835 | 0.145 | 0.933 | 0.121 | 0.315 |
| N-acetylornithine | Amino acid | 6205-08-9; | 0.99 | 0.79 | 0.80 | 0.992 | 0.994 | 0.368 | 0.933 | 0.145 | 0.331 |
| trans-4-hydroxyproline | Amino acid | 51-35-4; | 1.10 | 0.83 | 0.76 | 0.570 | 0.994 | 0.901 | 0.996 | 0.651 | 0.546 |
| creatine | Amino acid | 57-00-1; | 0.99 | 1.04 | 1.05 | 0.678 | 0.994 | 0.381 | 0.933 | 0.328 | 0.459 |
| creatinine | Amino acid | 60-27-5; | 0.96 | 0.85 | 0.88 | 0.716 | 0.994 | 0.619 | 0.974 | 0.877 | 0.619 |
| 5-methylthioadenosine (MTA) | Amino acid | 2457-80-9; | 0.93 | 0.97 | 1.04 | 0.630 | 0.994 | 0.808 | 0.996 | 0.084 | 0.315 |
| putrescine | Amino acid | 110-60-1; | 0.75 | 0.66 | 0.88 | 0.140 | 0.835 | 0.085 | 0.846 | 0.563 | 0.526 |
| spermidine | Amino acid | 124-20-9; | 0.95 | 0.92 | 0.97 | 0.468 | 0.994 | 0.322 | 0.933 | 0.069 | 0.299 |
| spermine | Amino acid | 71-44-3; | 0.98 | 1.05 | 1.07 | 0.959 | 0.994 | 0.522 | 0.961 | 0.600 | 0.529 |
| glutathione, reduced (GSH) | Amino acid | 70-18-8; | 1.01 | 0.90 | 0.89 | 0.975 | 0.994 | 0.349 | 0.933 | 0.070 | 0.299 |
| S-methylglutathione | Amino acid | 2922-56-7; | 0.90 | 0.74 | 0.82 | 0.208 | 0.835 | 0.120 | 0.933 | 0.124 | 0.315 |
| glutathione, oxidized (GSSG) | Amino acid | 103239-24-3; | 1.03 | 0.96 | 0.93 | 0.901 | 0.994 | 0.852 | 0.996 | 0.602 | 0.529 |
| ophthalmate | Amino acid | 495-27-2; | 0.87 | 0.65 | 0.74 | 0.681 | 0.994 | 0.198 | 0.933 | 0.003 | 0.085 |
| mannose-6-phosphate | Carbohydrate | 70442-25-0;104872-94-8; | 0.99 | 1.06 | 1.07 | 0.929 | 0.994 | 0.187 | 0.933 | 0.423 | 0.493 |
| sorbitol | Carbohydrate | 6706-59-8; | 0.65 | 0.80 | 1.23 | 0.047 | 0.591 | 0.407 | 0.933 | 0.562 | 0.526 |
| glucose-6-phosphate (G6P) | Carbohydrate | 103192-55-8; | 1.49 | 2.18 | 1.46 | 0.046 | 0.591 | 0.051 | 0.798 | 0.229 | 0.391 |
| glucose | Carbohydrate | 50-99-7; | 1.01 | 1.49 | 1.47 | 0.858 | 0.994 | 0.234 | 0.933 | 0.103 | 0.315 |
| Isobar: fructose 1,6-diphosphate, glucose 1,6-diphosphate | Carbohydrate |  | 1.00 | 1.41 | 1.41 | 0.943 | 0.994 | 0.547 | 0.961 | 0.103 | 0.315 |
| 3-phosphoglycerate | Carbohydrate | 80731-10-8; | 1.14 | 1.42 | 1.25 | 0.733 | 0.994 | 0.596 | 0.961 | 0.540 | 0.516 |
| lactate | Carbohydrate | 79-33-4; | 1.35 | 1.43 | 1.07 | 0.041 | 0.591 | 0.075 | 0.810 | 0.538 | 0.516 |
| citrate | Energy | 77-92-9; | 1.00 | 0.83 | 0.83 | 0.864 | 0.994 | 0.250 | 0.933 | 0.449 | 0.509 |
| fumarate | Energy | 100-17-8; | 1.12 | 0.89 | 0.79 | 0.590 | 0.994 | 0.724 | 0.996 | 0.505 | 0.516 |
| malate | Energy | 6915-15-7; | 1.02 | 1.09 | 1.08 | 0.988 | 0.994 | 0.750 | 0.996 | 0.600 | 0.529 |
| acetylphosphate | Energy | 94249-01-1; | 1.12 | 1.02 | 0.90 | 0.210 | 0.835 | 0.996 | 0.997 | 0.532 | 0.516 |
| phosphate | Energy | 7664-38-2 ; | 1.02 | 1.02 | 1.00 | 0.722 | 0.994 | 0.458 | 0.933 | 1.000 | 0.656 |
| pyrophosphate (PPi) | Energy | 1466-09-3; | 2.33 | 1.79 | 0.78 | 0.195 | 0.835 | 0.242 | 0.933 | 0.883 | 0.619 |
| linolenate [alpha or gamma; (18:3n3 or 6)] | Lipid |  | 0.94 | 0.88 | 0.93 | 0.609 | 0.994 | 0.519 | 0.961 | 0.794 | 0.599 |
| dihomo-linolenate (20:3n3 or n6) | Lipid |  | 1.47 | 1.43 | 0.97 | 0.179 | 0.835 | 0.455 | 0.933 | 0.798 | 0.599 |
| docosapentaenoate (n3 DPA; 22:5n3) | Lipid | 2234-74-4 ; | 1.11 | 1.41 | 1.26 | 0.643 | 0.994 | 0.279 | 0.933 | 0.184 | 0.358 |
| docosahexaenoate (DHA; 22:6n3) | Lipid | 6217-54-5; | 1.30 | 1.33 | 1.02 | 0.742 | 0.994 | 0.698 | 0.996 | 0.765 | 0.599 |
| caprylate (8:0) | Lipid | 124-07-2; | 1.04 | 0.78 | 0.75 | 0.423 | 0.994 | 0.423 | 0.933 | 0.423 | 0.493 |
| pelargonate (9:0) | Lipid | 112-05-0; | 1.00 | 1.05 | 1.05 | 0.975 | 0.994 | 0.742 | 0.996 | 0.677 | 0.563 |
| caprate (10:0) | Lipid | 112-37-8; | 0.99 | 1.04 | 1.05 | 0.930 | 0.994 | 0.829 | 0.996 | 0.891 | 0.619 |
| undecanoate (11:0) | Lipid | 334-48-5; | 0.99 | 1.15 | 1.16 | 0.792 | 0.994 | 0.787 | 0.996 | 0.289 | 0.431 |
| laurate (12:0) | Lipid | 143-07-7; | 1.03 | 1.01 | 0.98 | 0.343 | 0.994 | 0.941 | 0.996 | 0.904 | 0.619 |
| myristate (14:0) | Lipid | 544-63-8; | 1.09 | 1.09 | 1.01 | 0.643 | 0.994 | 0.588 | 0.961 | 0.906 | 0.619 |
| myristoleate (14:1n5) | Lipid | 544-64-9 ; | 1.06 | 1.05 | 0.98 | 0.908 | 0.994 | 0.958 | 0.996 | 0.865 | 0.619 |
| palmitate (16:0) | Lipid | 57-10-3; | 1.02 | 1.02 | 1.01 | 0.910 | 0.994 | 0.869 | 0.996 | 0.798 | 0.599 |
| palmitoleate (16:1n7) | Lipid | 373-49-9; | 0.75 | 0.95 | **1.28** | 0.475 | 0.994 | 0.875 | 0.996 | 0.020 | 0.225 |
| margarate (17:0) | Lipid | 506-12-7; | 1.14 | 0.98 | 0.86 | 0.160 | 0.835 | 0.860 | 0.996 | 0.307 | 0.445 |
| oleate (18:1n9) | Lipid | 112-80-1; | 1.22 | 0.92 | 0.75 | 0.053 | 0.591 | 0.846 | 0.996 | 0.341 | 0.462 |
| cis-vaccenate (18:1n7) | Lipid | 693-72-1; | 1.14 | 0.87 | 0.76 | 0.701 | 0.994 | 0.410 | 0.933 | 0.130 | 0.320 |
| linoleate (18:2n6) | Lipid | 60-33-3; | 1.14 | 1.14 | 1.00 | 0.874 | 0.994 | 0.864 | 0.996 | 0.968 | 0.645 |
| eicosenoate (20:1n9 or 11) | Lipid |  | 0.89 | 0.70 | 0.78 | 0.651 | 0.994 | 0.354 | 0.933 | 0.150 | 0.332 |
| dihomo-linoleate (20:2n6) | Lipid | 2091-39-6; | 1.23 | 0.94 | 0.76 | 0.514 | 0.994 | 0.784 | 0.996 | 0.197 | 0.368 |
| mead acid (20:3n9) | Lipid | 20590-32-3; | 0.86 | 0.76 | 0.88 | 0.673 | 0.994 | 0.595 | 0.961 | 0.374 | 0.486 |
| arachidonate (20:4n6) | Lipid | 506-32-1; | 1.11 | 1.06 | 0.95 | 0.924 | 0.994 | 0.997 | 0.997 | 0.610 | 0.529 |
| docosadienoate (22:2n6) | Lipid | 7370-49-2 ; | 1.08 | 0.82 | 0.76 | 0.833 | 0.994 | 0.517 | 0.961 | 0.195 | 0.368 |
| adrenate (22:4n6) | Lipid | 2091-25-0; | 1.15 | 2.03 | 1.76 | 0.980 | 0.994 | 0.453 | 0.933 | 0.074 | 0.299 |
| propionylcarnitine | Lipid | 17298-37-2 ; | 1.02 | 1.34 | 1.32 | 0.846 | 0.994 | 0.446 | 0.933 | 0.165 | 0.346 |
| butyrylcarnitine | Lipid | 25576-40-3; | 0.97 | 0.95 | 0.98 | 0.669 | 0.994 | 0.615 | 0.974 | 0.479 | 0.509 |
| carnitine | Lipid | 461-05-2; | 1.12 | 1.27 | 1.14 | 0.337 | 0.994 | 0.197 | 0.933 | 0.075 | 0.299 |
| 3-dehydrocarnitine* | Lipid | 10457-99-5; | 1.10 | 1.24 | **1.14** | 0.492 | 0.994 | 0.181 | 0.933 | 0.037 | 0.225 |
| acetylcarnitine | Lipid | 5080-50-2; | 1.19 | 1.27 | 1.07 | 0.345 | 0.994 | 0.270 | 0.933 | 0.400 | 0.493 |
| choline phosphate | Lipid | 72556-74-2; | 0.98 | **0.87** | **0.88** | 0.581 | 0.994 | <<br>0.001 | 0.096 | 0.040 | 0.225 |
| phosphoethanolamine | Lipid | 1071-23-4; | 1.08 | 1.03 | 0.96 | 0.430 | 0.994 | 0.843 | 0.996 | 0.857 | 0.619 |
| choline | Lipid | 67-48-1; | 0.98 | 1.05 | 1.07 | 0.760 | 0.994 | 0.980 | 0.997 | 0.292 | 0.431 |
| glycerol 3-phosphate (G3P) | Lipid | 29849-82-9; | 1.67 | 2.00 | 1.19 | 0.079 | 0.617 | 0.125 | 0.933 | 0.247 | 0.391 |
| glycerophosphorylcholine (GPC) | Lipid | 28319-77-9; | 1.43 | **1.97** | **1.38** | 0.070 | 0.606 | 0.029 | 0.565 | 0.002 | 0.085 |
| myo-inositol | Lipid | 87-89-8; | **0.75** | 0.89 | 1.18 | 0.010 | 0.591 | 0.587 | 0.961 | 0.382 | 0.486 |
| inositol 1-phosphate (I1P) | Lipid | 106032-59-1; | 0.96 | 0.97 | 1.00 | 0.653 | 0.994 | 0.988 | 0.997 | 0.855 | 0.619 |
| scyllo-inositol | Lipid | 488-59-5; | 1.27 | 1.10 | 0.87 | 0.631 | 0.994 | 0.423 | 0.933 | 0.698 | 0.575 |
| 2-palmitoylglycerophosphoethanolamine* | Lipid |  | 0.93 | 0.79 | 0.85 | 0.707 | 0.994 | 0.410 | 0.933 | 0.116 | 0.315 |
| 2-oleoylglycerophosphoethanolamine* | Lipid |  | 1.08 | 0.65 | **0.61** | 0.696 | 0.994 | 0.320 | 0.933 | 0.027 | 0.225 |
| 2-arachidonoylglycerophosphoethanolamine* | Lipid |  | 1.20 | 0.84 | 0.69 | 0.429 | 0.994 | 0.656 | 0.996 | 0.161 | 0.346 |
| 2-docosahexaenoylglycerophosphoethanolamine* | Lipid |  | 1.20 | 0.83 | 0.69 | 0.571 | 0.994 | 0.563 | 0.961 | 0.102 | 0.315 |
| 2-palmitoylglycerophosphocholine* | Lipid |  | **0.78** | 0.78 | 1.00 | 0.042 | 0.591 | 0.475 | 0.942 | 0.919 | 0.622 |
| 1-palmitoleoylglycerophosph | Lipid |  | 1.11 | 0.77 | 0.69 | 0.671 | 0.994 | 0.447 | 0.933 | 0.249 | 0.391 |
| ocholine* |  |  |  |  |  |  |  |  |  |  |  |
| 2-stearoylglycerophosphocholine* | Lipid |  | 0.58 | 1.36 | 2.36 | 0.346 | 0.994 | 0.164 | 0.933 | 0.135 | 0.325 |
| 2-oleoylglycerophosphocholine* | Lipid |  | 0.80 | 0.65 | 0.81 | 0.500 | 0.994 | 0.351 | 0.933 | 0.249 | 0.391 |
| 1-palmitoylglycerol (1-monopalmitin) | Lipid | 542-44-9; | 0.94 | 0.99 | 1.06 | 0.685 | 0.994 | 0.869 | 0.996 | 0.787 | 0.599 |
| 2-palmitoylglycerol (2-monopalmitin) | Lipid | 23470-00-0; | 1.05 | 1.15 | 1.09 | 0.295 | 0.994 | 0.249 | 0.933 | 0.468 | 0.509 |
| 1-stearoylglycerol (1-monostearin) | Lipid | 123-94-4; | 1.01 | 1.17 | 1.15 | 0.942 | 0.994 | 0.202 | 0.933 | 0.203 | 0.371 |
| sphingosine | Lipid | 123-78-4; | 1.08 | 0.98 | 0.91 | 0.798 | 0.994 | 0.860 | 0.996 | 0.403 | 0.493 |
| cholesterol | Lipid | 57-88-5; | 1.14 | 1.01 | 0.89 | 0.055 | 0.591 | 0.949 | 0.996 | 0.455 | 0.509 |
| hypoxanthine | Nucleotide | 68-94-0; | 1.08 | 1.10 | 1.03 | 0.885 | 0.994 | 0.834 | 0.996 | 0.793 | 0.599 |
| inosine | Nucleotide | 58-63-9; | 1.20 | 1.19 | 1.00 | 0.773 | 0.994 | 0.803 | 0.996 | 0.887 | 0.619 |
| adenine | Nucleotide | 73-24-5; | 0.89 | 0.73 | 0.81 | 0.360 | 0.994 | 0.187 | 0.933 | 0.256 | 0.391 |
| adenosine | Nucleotide | 58-61-7; | 1.14 | 1.20 | 1.06 | 0.158 | 0.835 | 0.139 | 0.933 | 0.644 | 0.546 |
| N1-methyladenosine | Nucleotide | 15763-06-1; | 1.09 | 1.19 | 1.09 | 0.772 | 0.994 | 0.557 | 0.961 | 0.593 | 0.529 |
| adenosine 5'-monophosphate (AMP) | Nucleotide | 149022-20-8; | 0.97 | 0.94 | 0.97 | 0.888 | 0.994 | 0.741 | 0.996 | 0.250 | 0.391 |
| adenosine 5'-diphosphate (ADP) | Nucleotide | 20398-34-9; | 0.77 | 0.94 | 1.22 | 0.060 | 0.591 | 0.525 | 0.961 | 0.092 | 0.315 |
| guanine | Nucleotide | 73-40-5; | 1.08 | 1.15 | 1.07 | 0.778 | 0.994 | 0.640 | 0.995 | 0.460 | 0.509 |
| guanosine | Nucleotide | 118-00-3; | 1.28 | 1.14 | 0.88 | 0.260 | 0.933 | 0.765 | 0.996 | 0.541 | 0.516 |
| guanosine 5'-monophosphate (GMP) | Nucleotide | 5550-12-9; | 0.90 | 0.86 | 0.96 | 0.694 | 0.994 | 0.522 | 0.961 | 0.532 | 0.516 |
| cytidine | Nucleotide | 65-46-3; | 0.96 | 0.80 | 0.83 | 0.749 | 0.994 | 0.440 | 0.933 | 0.169 | 0.346 |
| cytidine 5'-monophosphate (5'-CMP) | Nucleotide | 63-37-6; | 0.89 | 0.86 | 0.97 | 0.459 | 0.994 | 0.413 | 0.933 | 0.600 | 0.529 |
| thymidine 5'-monophosphate | Nucleotide | 33430-62-5; | 0.88 | 0.75 | 0.86 | 0.530 | 0.994 | 0.318 | 0.933 | 0.176 | 0.351 |
| uracil | Nucleotide | 66-22-8; | 1.11 | 0.87 | 0.79 | 0.866 | 0.994 | 0.450 | 0.933 | 0.458 | 0.509 |
| uridine | Nucleotide | 58-96-8; | 1.15 | 1.62 | 1.42 | 0.572 | 0.994 | 0.199 | 0.933 | 0.050 | 0.263 |
| methylphosphate | Nucleotide | 7023-27-0; | 1.10 | 1.54 | 1.40 | 0.350 | 0.994 | 0.118 | 0.933 | 0.114 | 0.315 |
| gulono-1,4-lactone | Cofactors and vitamins | 1128-23-0; | 0.71 | 0.61 | 0.86 | 0.083 | 0.617 | 0.532 | 0.961 | 0.940 | 0.631 |
| ascorbate (Vitamin C) | Cofactors and vitamins | 134-03-2; | 1.15 | 1.10 | 0.96 | 0.143 | 0.835 | 0.566 | 0.961 | 0.796 | 0.599 |
| nicotinamide | Cofactors and vitamins | 98-92-0; | 0.97 | 1.02 | 1.05 | 0.868 | 0.994 | 0.903 | 0.996 | 0.641 | 0.546 |
| nicotinamide adenine dinucleotide (NAD+) | Cofactors and vitamins | 53-84-9; | 0.85 | 0.65 | 0.76 | 0.344 | 0.994 | 0.031 | 0.565 | 0.028 | 0.225 |
| adenosine 5'diphosphoribose | Cofactors and vitamins | 68414-18-6; | 1.14 | 1.34 | 1.18 | 0.497 | 0.994 | 0.362 | 0.933 | 0.378 | 0.486 |
| pantothenate | Cofactors and vitamins | 137-08-6; | 0.85 | 0.78 | 0.91 | 0.437 | 0.994 | 0.343 | 0.933 | 0.251 | 0.391 |
| coenzyme A | Cofactors and vitamins | 85-61-0;18439-24-2; | 1.01 | 0.86 | 0.85 | 0.994 | 0.994 | 0.378 | 0.933 | 0.005 | 0.101 |
| acetyl CoA | Cofactors and vitamins | 102029-73-2; | 0.79 | 0.74 | 0.93 | 0.382 | 0.994 | 0.411 | 0.933 | 0.517 | 0.516 |
| alpha-tocopherol | Cofactors and vitamins | 59-02-9;10191-41-0; | 0.80 | 0.72 | 0.91 | 0.065 | 0.601 | 0.236 | 0.933 | 0.874 | 0.619 |

**Supplemental Table 2, related to Figure 1.** Expression Fold change of Glycolysis genes along the Paraxial mesoderm in a 9.5 day mouse embryo Fold changes of expression levels of Glycolysis genes shown in Figure 1H along the Paraxial mesoderm in 9.5 day old mouse embryo. Expression fold changes along the paraxial mesoderm domains correspond to ratios of normalized MAS values. Posterior Domain corresponds to the average of TB1, PSM1, PSM2 fragments while the Anterior Domain corresponds to an average of the fragments PSM3, PSM4, PSM5. Tailbud corresponds to TB1 fragments. A-PSM corresponds to PSM4, PSM5 fragments. Statistical significance was determined with an unpaired Student-test with the following cutoffs: ns p>0.05, *p<0.05, **p<0.01, ***p<0.001,****p<0.0001.

| Gene symbol | Affymetrix Probe ID | Posterior /Anterior |  | Tail bud/A-PSM |  |
| --- | --- | --- | --- | --- | --- |
|  |  | Fold change | P-value | Fold change | P-value |
| *Slc2a3* | 1437052_s_at | 3.59 | ** | 7.55 | ** |
| *Slc2a3* | 1455898_x_at | 4.95 | * | 11.02 | ** |
| *Pgm1* | 1453283_at | 1.91 | *** | 2.59 | **** |
| *Pgm2* | 1451149_at | 1.39 | ns | 2.06 | ** |
| *HK1* | 1437974_a_at | 1.20 | ns | 1.31 | ns |
| *Aldoa* | 1433604_x_at | 1.12 | ns | 1.39 | *** |
| *Aldoa* | 1434799_x_at | 1.09 | ns | 1.30 | ** |
| *Tpi1* | 1415918_a_at | 0.99 | ns | 1.14 | ns |
| *Bpgm* | 1415864_at | 2.05 | ** | 3.25 | *** |
| *Bpgm* | 1415865_s_at | 1.93 | *** | 2.87 | ** |
| *Eno3* | 1417951_at | 1.69 | * | 1.27 | ** |
| *LDHB* | 1448237_x_at | 1.30 | * | 1.39 | ns |
| *LDHB* | 1455235_x_at | 1.32 | * | 1.38 | ns |

**Supplemental Table 3, related to figure 2.** Expression Fold change of Glycolysis genes along the Paraxial mesoderm in a 2-day old chicken embryo Fold changes of expression levels of Glycolysis genes along the Paraxial mesoderm in Stage 12HH chicken embryo. Expression fold changes along Paraxial mesoderm domains corresponds to ratios of normalized MAS values. Posterior Domain corresponds to the average of (TB1, PSM1, PSM2, PSM3) while the Anterior Domain corresponds to an average of fragments (PSM4, PSM5, PSM6, PSM7, PSM8). Tailbud corresponds to (TB1) fragments. A-PSM corresponds to (PSM6, PSM7, PSM8). Statistical significance was determined with an unpaired Student-test with the following cutoffs: ns p>0.05, *p<0.05, **p<0.01, ***p<0.001,****p<0.0001.

| Gene symbol | Affymetrix Probe ID | Posterior/Anterior |  | Tail bud/A-PSM |  |
| --- | --- | --- | --- | --- | --- |
|  |  | Fold change | P-value | Fold change | P-value |
| *SLC2A1* | Gga.1040.1.S1_at | 1.66 | ** | 1.25 | ns |
| *HK1* | Gga.2197.1.S1_s_at | 1.37 | ** | 1.50 | ns |
| *Gpi1* | GgaAffx.20134.1.S1_s_at | 1.97 | ** | 2.57 | ** |
| *PFKFB4* | GgaAffx.25392.2.S1_s_at | 1.36 | ns | 1.59 | ns |
| *PFKP* | Gga.5753.1.S1_s_at | 1.35 | ** | 1.61 | * |
| *G3P* | Gga.1374.4.S1_x_at | 1.09 | * | 1.18 | * |
| *PGK1* | Gga.4503.1.S1_a_at | 1.45 | ** | 1.74 | * |
| *PGAM1* | Gga.6033.1.S1_s_at | 1.33 | * | 1.35 | ns |
| *ENOA* | Gga.1383.1.S1_at | 1.14 | ** | 1.24 | * |
| *PKM* | Gga.4299.1.S1_at | 1.34 | *** | 1.45 | * |
| *LDHB* | Gga.4149.1.S1_at | 1.42 | **** | 1.66 | *** |

## Supplementary Movie Legends

**Supplementary movie 1:** Elongation of the embryo in control, 2DG and NaN3 conditions. The starting time of the movie corresponds to the time when the analysis was started (corresponding to Figure 3 D). Bright field (left) and corresponding cell trajectories of PSM cells electroporated with an H2B-Venus (nuclear) construct (right) are shown for each condition. Only a portion of the trajectories are shown for clarity. The starting time of the movie corresponds to the time when the analysis was started (corresponding to Figure 3 DF).

**Supplementary movie 2:** Bright field (left) and corresponding cell trajectories of PSM cells electroporated with an H2B-Venus (nuclear) construct (right) are shown for each condition. Only a portion of the trajectories are shown for clarity. The starting time of the movie corresponds to the time when the analysis was started (corresponding to Figure 3 G-

**Supplementary movie 3:** Elongation of the embryo in control and alkaline conditions. Bright field (left) and corresponding cell trajectories of PSM cells electroporated with an H2B-Venus construct (right) are shown for each condition. The starting time of the movie corresponds to the time when the analysis was started (corresponding to Figure 3 (P-R)).

**Supplementary Movie 4:** Effect of 2DG on the differentiation of the neuro-mesodermal precursors (NMPs). H2B-RFP was electroporated in the anterior primitive streak area containing the NMPs of stage 4-5HH control (left) and 2DG-treated chicken embryos (right). In the control embryo, electroporated NMPs produce descendants in the neural tube and in the paraxial mesoderm. In contrast, 2DG treatment inhibits the production of paraxial mesoderm cells while NMP descendants accumulate in the Neural Tube.

## Supplemental experimental procedures

### Mouse embryos dissection and metabolome analysis

The posterior end of three hundred day 9.5 CD1 mouse embryos was dissected into three consecutive fragments of roughly equivalent size along the antero-posterior axis as shown in Figure 1A. The most posterior fragment includes the tail bud and the level of the posterior PSM, the next level correspond to the level of the anterior PSM and the most anterior level corresponds to the most posterior somitic region. Embryos were dissected in cold PBS on silicone-covered Petri dishes and frozen by batches of five in liquid nitrogen. Triplicate pools of fragments of each level were created. Samples were then shipped to Metabolon Inc. (Durham, NC, USA) for metabolomics analysis.

#### Sample Preparation

Each sample is accessioned into a LIMS system, assigned a unique identifier, and stored at −70°C. To remove protein, dissociate small molecules bound to protein or trapped in the precipitated protein matrix, and to recover chemically diverse metabolites, proteins are precipitated with methanol, with vigorous shaking for 2 minutes (Glen Mills Genogrinder 2000). The sample is then centrifuged, supernatant removed (MicroLab STAR^®^ robotics), and split into equal volumes for analysis on the LC+, LC-, and GC platforms; one aliquot is retained for backup analysis, if needed.

#### Liquid Chromatography/Mass Spectrometry (LC/MS/MS) and Gas Chromatography/Mass Spectrometry (GC/MS)

The LC/MS portion of the platform incorporates a Waters Acquity UPLC system and a Thermo-Finnigan LTQ mass spectrometer, including an electrospray ionization (ESI) source and linear ion-trap (LIT) mass analyzer. Aliquots of the vacuum-dried sample are reconstituted, one each in acidic or basic LC-compatible solvents containing 8 or more injection standards at fixed concentrations (to both ensure injection and chromatographic consistency). Extracts are loaded onto columns (Waters UPLC BEH C18-2.1 × 100 mm, 1.7 μm) and gradient-eluted with water and 95% methanol containing 0.1% formic acid (acidic extracts) or 6.5 mM ammonium bicarbonate (basic extracts). Samples for GC/MS analysis are dried under vacuum desiccation for a minimum of 18 hours prior to being derivatized under nitrogen using bistrimethyl-silyl-trifluoroacetamide (BSTFA). The GC column is 5% phenyl dimethyl silicone and the temperature ramp is from 60° to 340° C in a 17 minute period. All samples are then analyzed on a Thermo-Finnigan Trace DSQ fast-scanning single-quadrupole mass spectrometer using electron impact ionization. The instrument is tuned and calibrated for mass resolution and mass accuracy daily.

#### Quality Control

All columns and reagents are purchased in bulk from a single lot to complete all related experiments. For monitoring of data quality and process variation, multiple replicates of a pool of human plasma are injected throughout the run, interspersed among the experimental samples in order to serve as technical replicates for calculation of precision. In addition, process blanks and other quality control samples are spaced evenly among the injections for each day, and all experimental samples are randomly distributed throughout each day’s run. In our preliminary human plasma sample analysis, median relative standard deviation (RSD) was 13% for technical replicates and 9% for internal standards.

#### Bioinformatics

The LIMS system encompasses sample accessioning, preparation, instrument analysis and reporting, and advanced data analysis. Additional informatics components include data extraction into a relational database and peak-identification software; proprietary data processing tools for QC and compound identification; and a collection of interpretation and visualization tools for use by data analysts. The hardware and software systems are built on a web-service platform utilizing Microsoft’s .NET technologies which run on high-performance application servers and fiber-channel storage arrays in clusters to provide active failover and load-balancing.

#### Compound Identification, Quantification, and Data Curation

Biochemicals are identified by comparison to library entries of purified standards. At present more than 2400 commercially available purified standards are registered into LIMS for distribution to both the LC and GC platforms for determination of their analytical characteristics. Chromatographic properties and mass spectra allow matching to the specific compound or an isobaric entity using proprietary visualization and interpretation software. Additional recurring entities may be identified as needed via acquisition of a matching purified standard or by classical structural analysis. Peaks are quantified using area under the curve. Subsequent QC and curation processes are designed to ensure accurate, consistent identification, and to minimize system artifacts, mis-assignments, and background noise. Library matches for each compound are verified for each sample.

#### Statistical Analysis

Missing values (if any) are assumed to be below the level of detection. Given the multiple comparisons inherent in analysis of metabolites, between-group relative differences are assessed using both Student’s t-tests (p-value) and false discovery rate analysis (q-value). Pathways are assigned for each metabolite, also allowing examination of overrepresented pathways. Hierarchical clustering analysis was performed using Pearson correlation coefficient with the MeV 4.9 (TM4) software.

### Chicken embryo culture

Fertilized chicken eggs were obtained from commercial sources. Eggs were incubated at 38 °C in a humidified incubator, and embryos were staged according to Hamburger and Hamilton (HH) (*1*). In this study, we started to culture chicken embryos mainly from stage 9HH at 37°C using the Early Chick (EC) culture system (*2*). For drug treatments, 2mM 2DG (2-Deoxy-D-glucose; Sigma), 1mM NaN3 (sodium azide; Sigma), or albumin plates were prepared. For signaling inhibitors, 500μM SU5402 (Sigma), 50 μM PD035901 (AXON MEDCHEM BV), 100 μM DAPT (Sigma) or 50 μM BMS204493 (synthesized by Novalix) were diluted in PBS, and 100 μl of the inhibitor solution was added under and on top of the embryos. For alkaline plates, 23mM NaOH albumin plates (2 ml) were prepared, so that they reached pH11.0-11.3 (the pH of control albumin plates is 9.4-9.6).

### PSM microdissection and Microarray

PSM microdissection was performed as described (*3*). Stage 12HH chicken embryos were pinned on a silicon-coated petri dish in PBS. After removing the endoderm and the ectoderm, both the left and right posterior paraxial mesoderm from the same embryo were dissected into 10 pieces each. All the fragments were stored in Trizol (Invitrogen) at −80°C for subsequent RNA extraction. Biotinylated cRNA targets were prepared from total RNA using a double amplification protocol according to the GeneChip^®^ Expression Analysis Technical Manual: Two-Cycle Target Labeling Assay (P/N 701021 Rev.5, Affymetrix, Santa Clara, USA). Following fragmentation, cRNAs were hybridized on GeneChip^®^ chicken Genome arrays. Each microarray was then washed and stained on a GeneChip fluidics station 450 and scanned with a GeneChip Scanner 3000 7G. Finally, raw data (.CEL Intensity files) were extracted from the scanned images using the Affymetrix GeneChip Command Console (AGCC) version 3.1. CEL files were further processed with MAS5 and RMA algorithms using the Bioconductor package (version 2.8) available through R (version 2.12.1). Probe sets were filtered based on their expression intensity value (MAS5 value). Fold changes were compared between average of MAS values of each fragment for mouse triplicate microarray series (*3*) or chicken duplicated microarray series. Statistical comparisons were performed with unpaired two-tailed student t-test. Microarray data were deposited in GEO database under the accession number of GSE39613 for Mouse and GSE75798 for chicken samples.

### Sample preparation and measurement of metabolic activity

The posterior end of day 9.5 CD1 mouse embryos and stage 11 HH chicken embryos were dissected using tungsten needles in cold PBS into 3 parts corresponding to the levels of the posterior PSM (P-PSM), the anterior PSM (A-PSM) and the newly formed somites (Figure1A). For each analysis, fragments of the same level from 3 embryos were immediately frozen in liquid nitrogen and pooled. Whole cell lysates were prepared by pipetting and vortexing in *PBS* containing 0.5% Tween 20 and protease inhibitor cocktails (Roche). Each sample was normalized by measuring total protein levels using Bio-Rad protein assay kit (Bio-Rad). To analyze the effect of drugs on metabolic activity, stage 9 HH chicken embryos were incubated at 37°C with inhibitors in EC culture as described above. After 10 h of incubation, the posterior end of treated embryos was dissected and samples were prepared as described above. Cellular lactate levels were measured using a Lactate assay kit (biovision) in accordance with the manufacturer’s instructions. Cellular ATP levels were measured using ATPlite Luminescence ATP Detection Assay System (Perkin-Elmer) in accordance with the manufacturer’s instructions. Cytochrome-C oxidase activity was measured using CytoCHROME C oxidase assay kit (Sigma) in accordance with the manufacturer’s instructions. Biological triplicates of the experiments were performed, and experiments were reproduced at least twice for each measurement of metabolic activity. Values obtained for these assays were normalized by P-PSM or by control values to compare the different AP levels.

### Time-lapse microscopy and axis elongation measurements

Stage 9HH chicken embryos were cultured ventral side up on a microscope stage using a custom built time-lapse station (*4*). We used a computer controlled, wide-field (10×objective) epifluorescent microscope (Leica DMR) workstation, equipped with a motorized stage and cooled digital camera (QImaging Retiga 1300i), to acquire 12-bit grayscale intensity images (492 × 652 pixels). For each embryo, several images corresponding to different focal planes and different fields were captured at each single time-point (frame). The acquisition rate used was 10 frames per hour (6 min between frames). To quantify axis elongation length, the image sequence was first registered to the last formed somite at the beginning of the time-lapse experiment (yellow asterisk in Fig 3), and the advancement of the Hensen’s Node was tracked as a function of time using the manual tracking plug-in in Image J (*5*).

### Cell trajectory analysis

To study the motility of cells in the presomitic mesoderm, we analyzed cell trajectories in the posterior region of the PSM in an area of about 400 μm representing 1/3 of the PSM length, as shown schematically with a box in Figure 3 (G-I). Because of the small depth of the PSM with respect to its area, the motion of the cells can be considered two dimensional. From the trajectories of cells we obtain their time-averaged displacement, or the mean square displacement, given by: MSD = 〈Δ*r*^2^ (*t*)〉, where 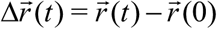 defines the distance that the cell travels in a time *t*, known as the lag time. When the diffusive motion in coupled with a drifting flow, the MSD is given by

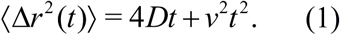

where *D* is the diffusion coefficient and *v* is the drift, related to the overall movement of the tissue (*6*). For each cell trajectory, the MSD is calculated and adjusted with Eq. (1) to obtain *D*. Cell tracking was performed on fluorescent images using the Mosaic plugin (MOSAIC Group, MPI-CBG, Dresden) for ImageJ (*7*). Further analyses to extract *D* were done using custom made Matlab (MathWorks) routines.

### Whole mount *in-situ* hybridization

Stage 9HH embryos were cultured with and without drugs at 38 °C in EC culture. After 16 h of incubation with 2DG, 10hr incubation with PD035901 embryos were fixed in 4% paraformaldehyde (PFA). Whole mount *in situ* hybridization was carried out as described(*8*). Probes for *FGF8* (*9*), *AXIN2* (*10*), *CMESO1* (*11*), *CMESPO* (*12*), *SPRY2*(*13*) and *BRACHYURY*(*14*) have been described. Probes for *PKM*, *LDHB* were generated from chicken embryo cDNA by PCR using published sequences.

### CMESPO Antibody generation

cDNA coding for the full length chicken CMESPO was cloned in pET vector expression system (Novagen), expressed in *E.coli*. The recombinant protein was purified with His- Bind Kit (Novagen) and used to immunize rabbits (Cocalico Biologicals, Inc.). Sera were collected, assayed and validated by immunohistochemistry and used as anti- CMESPO/MESOGENIN1 polyclonal antibody.

### Immunohistochemistry

For whole mount immuno-histochemistry, 9.5 day CD1 mouse embryos and stage 12HH chicken embryos were fixed in 4% paraformaldehyde (PFA) at 4 °C overnight. Embryos were incubated with an antibody against Glut3 (1/300, Abcam) at 4 °C overnight, and next with secondary antibodies conjugated with AlexaFluor (Molecular probes) at 4 °C overnight. Images were captured using a laser scanning confocal macroscope *(TCS LSI; Leica) for Glut3 staining*.

For histological analysis, stage 9HH chicken embryos cultured with or without 2DG for 16h were fixed in 4% PFA. Embryos were then embedded in OCT compound and frozen in liquid nitrogen. Frozen sections (20μm) were incubated overnight at 4 °C with the primary antibody, and after washing they were incubated overnight at 4 °C with the secondary antibody conjugated with AlexaFluor (Molecular probes). We used antibodies against T/BRACHYURY (1/1000, R&D Systems: AF2085), SOX2 (1/1000, Millipore: ab5603), CTNNB1/β-Catenin (1/500, BD transduction laboratories: #610153): For CTNNB1 staining, antigen retrieval (Incubation with Target Retrieval Solution Citrate pH6 (DAKO) at 105 °C for 10min) was needed before first antibody incubation. The BRACHYURY and SOX2 staining images were captured using a laser scanning confocal microscope with a 40X objective (TCS SP5; Leica). To image the whole PSM, we used the tiling and stitching function of the microscope (3 by 2 matrix). *For CMESPO and CTNNB1 stainings, images were captured using a laser scanning confocal microscope with a 20X objective (TCS SP5; Leica). Whole images were created by tiling the scans of 8 images. High magnification images were captured with a 63X objective*.

### Plasmid preparation and electroporation

pCAGGS-Venus and pCAGG-H2B-RFP have been described (*15*). Chicken embryos ranging from stage 6HH to stage 7HH were prepared for EC culture. A DNA solution (1.0-5.0 μg/μl) was microinjected in the space between the vitelline membrane and the epiblast surrounding the anterior primitive streak level which contains the precursors of the paraxial mesoderm. *In vitro* electroporations were carried out with five successive square pulses of 8V for 50ms, keeping 4mm distance between anode and cathode using Petri dish type electrodes (CUY701P2, Nepa Gene, Japan) and a CUY21 electroporator (Nepa Gene, Japan). This procedure only labels the superficial epiblast layer. For time-lapse analysis, after electroporation, embryos were re-incubated in a humidified incubator until they reached stage 9HH at 38 °C, then embryos were transferred to the microscope stage for time-lapse imaging.

### Cell proliferation and apoptosis analysis

Stage 9 HH chicken embryos were cultured on agar plates containing 2DG or NaN3 at 38 °C for 12 h. Embryos were fixed in 1% PFA at 4 °C for 20min. Whole mount TUNEL staining was performed using the ApopTag Red *In Situ* kit (#S7165; Millipore), and proliferating cells were stained using anti-Phospho-histone H3 (pH3) antibody (1/1000, Millipore). Images were captured using a laser scanning confocal microscope (TCS SP5; Leica). Ratios of proliferating and apoptotic cells were calculated by manual counting for DAPI, pH3 and TUNEL positive cells.

### Quantitative RT-PCR

Stage 9 HH embryos were cultured with or without 2mM 2DG and PD035901 at 38 °C for 10 h. Then total RNA was extracted from dissected tailbud regions. 500ng~1pg total RNA was used as template for cDNA synthesis using the QuantiTect kit (Qiagen) or Super script III (Thermo Fisher Scientific). RT-PCR was performed using QuantiFast SYBR Green RT-PCR Kit (Qiagen) primers or iTaq Universal SYBR Green Supermix (Bio-Rad) and run on a LightCycler 480II (Roche) or CFX384 Touch qPCR System (Bio-Rad). Beta-Actin was used as an internal control. Primers were as follows: *GLUT1* (SLC2A1-L: AGTACGGAGAGCATTCCCCT, SLC2A1-R: CTCAGGAAGGTGGGAAGCTG), HK1 (HK1-L: GAGTTCAAGCTCACCCACGA, HK1-R: CTTCTTCAGGCCTGCTTCCA), *G3P* (G3P-L: GAACTGAGCGGTGGTGAAGA, G3P- R: CCACATGGCATCCAAGGAGT), *T* (T-L: CGAGGAGATCACAGCTTTAAAAATT, T-R: TCATTTCTTTCCTTTGCGTCAA), *CMESPO* (CMESPO- L: AAAGCCAGTGAGAGGGAGAA, CMESPO-R: GGTGCACTTGAGGGTCTGTA), *SOX2* (SOX2-L: GCAGAGAAAAGGGAAAAAGGA, SOX2-R: TTTCCTAGGGAGGGGTATGAA), *SOX1* (SOX1-L: AGGAGAATCCCAAGATGCAC, SOX1-R: CTCCGACATCACCTTCCAC), *SAX1* (SAX1- L:CAGCTTTCACCTACGAGCAG, SAX1-R: TGGAACCAGATCTTCACCTG), *BETA-ACTIN* (Gg_ACTB_1_SG QuantiTect Primer Assay), GPI(Gg_GPI_1_SG QuantiTect Primer Assay), PFKP(Gg_PFKP_1_SG QuantiTect Primer Assay), PGK1(Gg_PGK1_1_SG QuantiTect Primer Assay), ENO1(Gg_ENO1_1_SG QuantiTect Primer Assay), PKM(Gg_PKM2_1_SG QuantiTect Primer Assay), *LDHB*( Gg_LDHB_1_SG QuantiTect Primer Assay), *AXIN2* (Gg_AXIN2_1_SG QuantiTect Primer Assay). Data were normalized by control samples.

### Glucose uptake

100μl 1mM 2-NDBG (Life technologies) was added both on top and below stage 11 to 12 HH chicken embryos and incubated at 38 °C for 2 h in EC culture. After culture, embryos were fixed in 4% PFA for 20 min, and then washed with PBS for 5 min at least 3 times. Images were captured using a laser scanning confocal microscope (TCS SP5; Leica) or macroscope (*TCS LSI; Leica*).

### Extracellular pH measurement

Stage 9HH embryos were incubated with and without 2DG in EC culture at 38 °C for 7 h. Next, embryos were transferred either to control plates (pH 9.6), 2DG-containing plates or alkaline plates (pH 11). Then 100μl 25μM pH sensitive dye (pHrodo Red intracellular pH Indicator: Lifetechnologies) was added both on top and below embryos, which were then incubated at 38 °C for 3 h. Embryos were then washed once in PBS, and mounted on MatTek glass-bottom dishes dorsal side up on a thin albumin agar layer (control, 2DG or Alkaline). Images were captured with a laser scanning confocal microscope (TCS SP5; Leica) at 37 °C in a humidified atmosphere. In this protocol, pH indicator was mainly trapped on extracellular matrix or cellular membrane regions, indicating that it measures the extracellular pH. After image capture, **z-stacks (.lif) of individual embryos were rendered in FluoRender (Wan et al., 2012) to create maximum projection images centered on the midline of each embryo (cropped to leave ~400μm on each side of the midline). Intensities were measured on these images using the region of interest (ROI) and plot z-profile functionalities in ImageJ. Unpaired t-tests were applied when comparing average intensities between different experimental conditions. Paired t-tests were applied when comparing between regions the embryo.**

### Image acquisition and processing

All fluorescent images were acquired on Leica SP5 or *LSI* systems. Images were processed with Imagej Fiji and Adobe Photoshop, and MAX projection images are shown in Figures. Whole mount in situ images were captured on Leica Z16 APOA systems and DFC 420C camera, and Images were processed with Adobe Photoshop. Scale bars were measured using Imagej Fiji for confocal images, and by referencing somite size of control embryos for *in situ* images.

### Statistical analysis

Statistical significance for the comparisons between two groups of data (SupplementalTable 1, 2, Figure 2G, 4X), were assessed with unpaired two-tailed student t-test. Statistical significance for multiple groups comparison, (Figure1 E-G, L-N, 2 D-F, 3 A-C, Supplemental Figure4), were performed with one-way ANOVA and Tukey multiple comparison tests using GraphPad 6 (Prism).

